# The spreading of facultative H3K9me3-heterochromatin drives congenital disease

**DOI:** 10.1101/2023.08.11.552907

**Authors:** Chen Wang, Wei Zhang, Xue-Lin Chen, Wen-Fei Wang, Yong-Hao Li, Jian-Feng Chang, Xiao-Bo Guo, Yuan-Ya Jing, Ya-Bin Li, Xin-Yi Lu, Yu-Tong Li, Kai Liu, Jian-Quan Ni, Fang-Lin Sun

## Abstract

Heterochromatin marked by trimethylated histone 3 at lysine 9 (H3K9me3) plays fundamental roles in reprogramming to direct cell fate determination in higher eukaryotes. However, the upstream factors that guide the establishment and spreading of H3K9me3-heterochromatin, leading to human developmental malformations, remain elusive. In this study, we found that Cdk13, a member of RNA polymerase II (RNAPII) kinase, suppresses congenital heart syndrome by preventing global facultative H3K9me3-heterochromatin spreading. Additionally, Cdk13 directs the phosphorylation of a large set of heterochromatin proteins at specific sites, which are required for the interaction between HP1 and histone H3K9 methyltransferases. Furthermore, we identified a compound, an inhibitor of heterochromatin regulators, that can alleviate syndromic heart defects in *Cdk*13-mutant mice through the inhibition of H3K9me3-heterochromatin spreading. In summary, this study reveals a novel role and mechanism of Cdk13-triggered facultative H3K9me3-heterochromatin spreading in human genetic disease and paves the way for the treatment of congenital heart syndrome.

---

Facultative heterochromatin, marked by trimethylated histone 3 lysine 9 (H3K9me3), plays essential roles in cell lineage specification and maintenance of cell identity by silencing undesired genes (Jenuwein and Allis 2001; Grewal and Jia 2007; Riddle and Elgin 2018; Moazed 2011; Allshire and Madhani 2018; Nicetto and Zaret 2019). The writers and readers of canonical heterochromatin are key factors in initiating the assembly and spreading of H3K9me3-heterochromatin. The histone methyltransferases Su(var)3-9/Suv39h1/Clr4 ‘write’ H3K9me3, and the heterochromatin protein HP1 serves as a ‘reader’ of H3K9me2/3 (Hall et al. 2002; Volpe et al. 2002; Becker et al. 2017; Nicetto et al. 2019). Additionally, nuclear RNAs, meiotic messenger RNA elimination, and RNA interference machinery are all critical regulators of H3K9me3-heterochromatin (Zofall et al. 2012; Hiriart et al. 2012; Lee et al. 2013; Yamanaka et al. 2013; Tashiro et al. 2013; Chalamcharla et al. 2015). The mechanism that guides the dynamics of facultative H3K9me3-heterochromatin spreading within gene-rich euchromatic regions, however, is still largely unknown.

In a previous screen aimed at identifying upstream regulators of heterochromatin formation and spreading (Pan et al. 2015), we discovered that dCdk12, a known kinase member that phosphorylates the C-terminal repeat domain of RNAPII (Even et al. 2006; Chen et al. 2007; Bartkowiak et al. 2010; Wu et al. 2018) controls the spreading of HP1 and H3K9me3 in the euchromatic domains of the *Drosophila melanogaster* genome. Cdk13 in mammals is one of the orthologues of dCdk12 (Dai et al. 2012; Bösken et al. 2014; Liang et al. 2015; Greifenberg et al. 2016). Mutations in *CDK*13 have been linked to syndromic congenital heart defects (Sifrim et al. 2016; Deciphering Developmental Disorders Study 2017) and other phenotypes, such as growth retardation, craniofacial abnormality and intellectual disorder in humans (Bostwick et al. 2017; Akker et al. 2017; Hamilton et al. 2019); however, the underlying mechanism of *CDK*13 mutation-resulted human genetic diseases remain elusive.

## Expansion of H3K9me3-marked heterochromatin after *Cdk*13 depletion

To investigate whether Cdk13 affects H3K9me3-heterochromatin formation and cellular programming in mammals, we first depleted *Cdk*13 from mouse embryonic fibroblasts (MEFs) by specific short hairpin RNA (shRNA) knockdown (KD) (Supplementary Fig. 1A). *Cdk*13 KD resulted in approximately a 50% reduction in the number of cells compared to the control cells after 3-5 days of culture, indicating that the proliferation of MEFs requires Cdk13 (Supplementary Fig. 1B). Interestingly, immunofluorescence staining experiments showed that the intensity of classical heterochromatin markers, such as H3K9me3, HP1α, HP1β, and Trim28/Kap1, was significantly elevated in *Cdk*13 KD MEFs compared with controls (Fig. 1A, Supplementary Fig. 1C,D). Markers of active chromatin, including H3K9ac, H3K27ac, and H4K16ac, were significantly reduced Supplementary Fig. 1E-G). The expansion of heterochromatinization upon *Cdk*13 KD was further confirmed by transmission electron microscopy (TEM), and a robust increase in darker or condensed heterochromatin was evident after *Cdk*13 depletion (Fig. 1b, Supplementary Fig. 2A). These findings support the notion that Cdk13 in mice, similar to its orthologue dCdk12 in fruit flies (Pan et al. 2015), plays a critical role in preventing the spreading of heterochromatin.

**Fig. 1.**
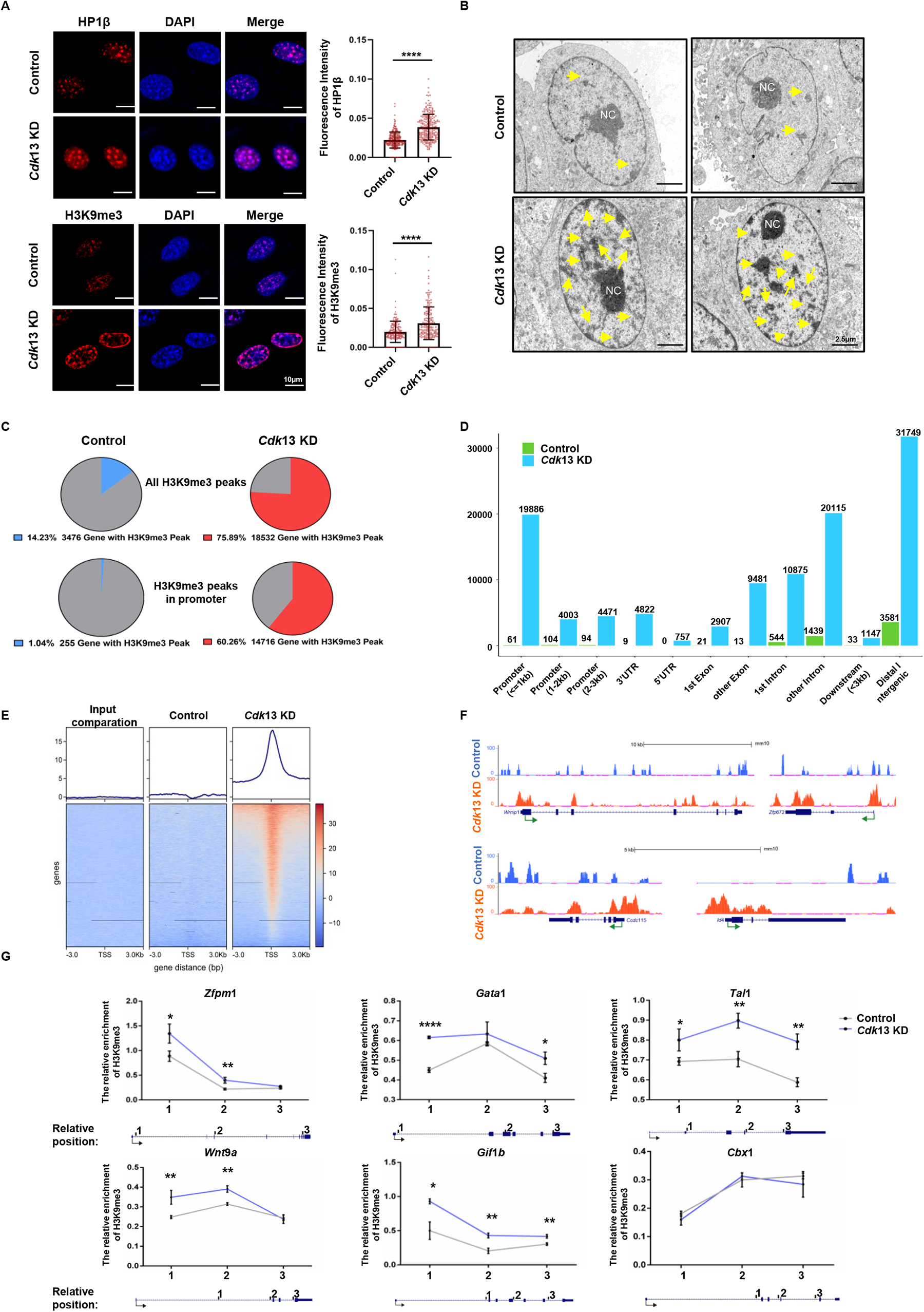
Global increase in H3K9me3-heterochromatinization after *Cdk*13 depletion. **A**, Left, Representative immunofluorescence staining for HP1β (red) and H3K9me3 (red) in control and *Cdk*13 knockdown (KD) mouse embryonic fibroblasts (MEFs) with lentivirus shRNA. Right, Quantification of fluorescence intensity of HP1β and H3K9me3. Each point represents a cell, and more than 300 cells per condition were analyzed. The data are presented as the means ± SEMs. \*\*\**P* < 0.001, \*\*\*\**P* < 0.0001, t test. **B**, Representative transmission electron microscopy images of *Cdk*13 KD MEFs and control MEFs are shown. Fifteen nuclei were analysed from each group, with arrow indicating the condensed heterochromatin. NC, nucleolus **C**, The H3K9me3 globally increased in *Cdk*13 KD MEFs compared to controls. The Pie plot shows the percentage of genes with H3K9me3 peaks; Above: The number and percentage of genes with H3K9me3 peaks among the known genes in the entire mouse genome; Below: The number and percentage of genes with H3K9me3 peaks in the promoter region (transcriptional start site ± 3 kb) among the H3K9me-enriched genes before (Control) and after *Cdk*13 KD. Blue represents the percentage in the control group, while red represents the percentage in the *Cdk*13 KD group. **D**, The enrichment of H3K9me3 peaks in all regions of euchromatic genes and intergenic regions. **E**, H3K9me3 dramatically increased in *Cdk*13 KD MEFs around transcriptional start sites (TSS). **F**, Representative genes with increased H3K9me3 after *Cdk*13 depletion are shown. The views of H3K9me3 in the UCSC genome are indicated. Blue represents control group, and Red represents *Cdk*13 KD group. TSS and the direction of transcription is indicated by arrows. **G**, ChIP-qPCR analysis confirmed the H3K9me3 enrichments in *Cdk*13 KD MEF cells compared to control cells (Control). The HP1β gene was used as a negative control. Three pairs of primers, within or adjacent to the gene exon, were designed for each gene.

To examine the precise genomic distribution of H3K9me3-heterochromatin upon depletion of *Cdk*13, we performed chromatin immunoprecipitation (ChIP) analysis using control and *Cdk*13-KD MEFs. Following ChIP sequencing, an increased association with H3K9me3 was observed in more than 18,500 known mouse genes, accounting for nearly 75.9% of known genes in the entire mouse genome (Fig. 1C). This indicates a general role of Cdk13 in global heterochromatin spreading at euchromatic regions. The most prominent increase in H3K9me3 peaks includes gene promoters, gene exons/introns, and distal intergenic regions (Fig. 1D), corresponding to 18.4%, 20.5%, and 28.8% of overall increased peaks in *Cdk*13 KD MEFs, respectively (Supplementary Fig. 2B). Further analysis of the increased H3K9me3 peaks distributed in the promoters, untranslated regions, and exons of target genes revealed more than 10-fold increase in H3K9me3 peaks compared to those in MEF control cells, calculated by the relative percentage of H3K9me3 peaks (Supplementary Fig. 2B,C). In contrast, no significant change was observed in the overall percentage of peaks in intronic regions upon *Cdk*13 KD (Supplementary Fig. 2B). Notably, the increase in H3K9me3 in the H3K9me3-enriched genes preferentially occurred at the region flanking the transcription start site (TSS) (Fig. 1E,F). While a subset of genes showed a similar increase in enrichments at both the TSS and gene body regions (Fig. 1F and Supplementary Fig. 2C).

The distal intergenic region remained to be the top H3K9me3-enriched region upon *Cdk*13 KD (Fig. 1D), however, exhibited a more than 50% relative reduction in percentage of H3K9me3 peaks, decreasing from 60.7% in control MEFs to 28.8% in *Cdk*13 KD MEFs (Supplementary Fig. 2B). The increased H3K9me3 in several euchromatic genes was confirmed by quantitative PCR using anti-H3K9me3 CHIP DNA of control and *Cdk*13 KD MEF cells (Fig. 1G). Overall, these results confirmed the general role of Cdk13 in preventing global H3K9me3-heterochromatin spreading at euchromatic loci, containing protein-coding genes with pleiotropic cellular functions including cardiomyocyte function, transcription regulation, and cell cycle regulation in the mouse genome (Supplementary Fig. 2D-G).

## *Cdk*13 mutant mice display heart defects, increased heterochromatinization and specific gene silencing in cardiomyocytes

To verify the role of Cdk13 in H3K9me3-heterochromatin observed in *in vitro* cellular experiments, we generated *Cdk*13-knockout (*Cdk*13*^-/-^*) mice (Supplementary Fig.3A,B). Most *Cdk*13^-/-^ mice died before E14 (Supplementary Table 1), supporting a critical role of Cdk13 in embryonic development. The ratio of surviving *Cdk*13^-/-^ embryos at E11.5 was only 16.9% of total progeny, which was much less than the expected 25%. By E13.5, the number of surviving *Cdk*13^-/-^ embryos dramatically decreased to 4.9% of the total progeny (Supplementary Table 1). Detailed analysis of the *Cdk*13^-/-^ embryos showed growth retardation as early as E9.5 and E11.5 (Supplementary Fig. 3C,D). Severe abnormalities were apparent in E13.5 *Cdk*13^-/-^ embryos, including craniofacial defects, digital anomalies, pleural/back effusion, and smaller body size (Supplementary Table 2, Supplementary Fig. 3E,F), which is consistent with a previous phenotypic report (Monika et al. 2019). Further dissection results showed that nearly 90% Cdk13^-/-^ mutant embryos displayed defective interventricular septum (Fig. 2A) and reduced thickness of the ventricular walls (Supplementary Fig. 4A, B).

**Fig. 2.**
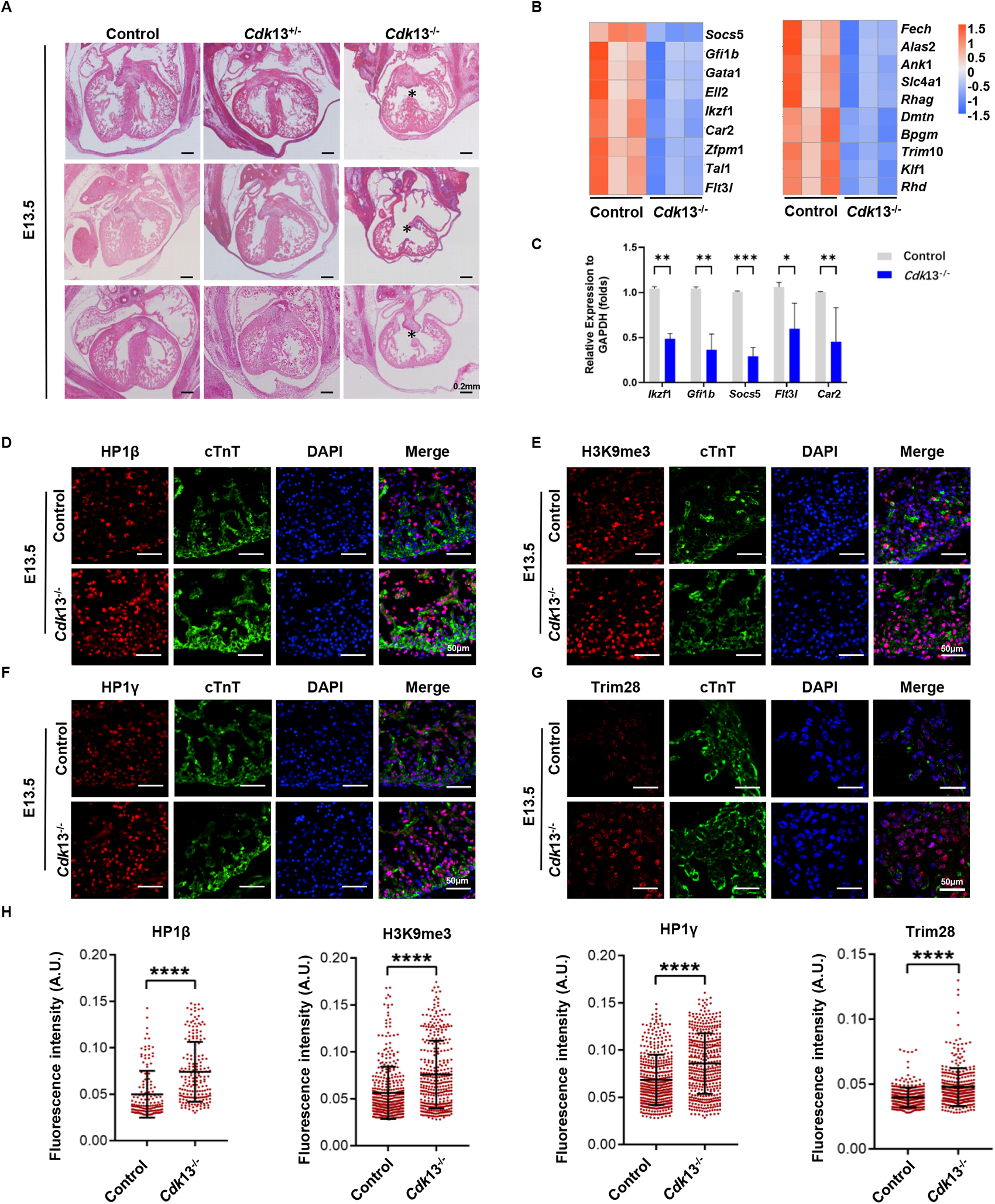
Deletion of *Cdk*13 in mice causes embryonic morphology defects and increased H3K9me3-heterochromatinization in cardiomyocytes. **A**, *Cdk*13^-/-^ resulted in abnormal cardiac structure such as lack of interventricular septum in mouse E13.5 embryos. The embryos were stained with haematoxylin and eosin. The left panel shows the control hearts, the middle panel shows *Cdk*13*^+/-^* embryo hearts, and the right panel shows *Cdk*13*^-/-^* embryo hearts. Asterisks indicate the area with ventricular septal defects. **B**, The heatmap shows downregulated genes related to development (left) and myeloid cells (right). **C**, qRT-PCR confirmed the reduced transcription level of randomly selected genes from (d) in *Cdk*13^-/-^ E13.5 heart tissue. The data are presented as the means ± SEMs. \**P* < 0.05, \*\**P*<0.01, \*\*\**P* < 0.001, t-test. **D-G**, Increased heterochromatinization in E13.5 *Cdk*13^-/-^ embryonic heart cells. Immunofluorescence staining was performed using antibodies against known markers of heterochromatin, including HP1β (**D)**, H3K9me3 (**E)**, HP1γ (**F)**, and Trim28 (**G)**, which are labelled in red. cTnT-labelled cardiomyocytes are shown in green, and DAPI is shown in blue. All images show the left ventricle position. The scale bar was 50 μm. **H**, The fluorescence intensity of single cells is shown according to the immunofluorescence staining results (**D-G**). Each point represents a cell, and more than 300 cells per condition were analyzed. A. U, arbitrary unit, *n* = 3, t-test. \*\*\*\**P* < 0.0001.

To examine the transcriptional role of Cdk13, we performed RNA-sequencing experiments. RNA-sequencing analysis of gene expression in surviving E10.5 *Cdk*13*^-/-^* mouse hearts revealed that the transcription of over 70% of affected genes was downregulated in *Cdk*13*^-/-^*hearts compared to controls (Fig. 2B, Supplementary Fig. 4C). The reduction in transcription was confirmed by qRT-PCR (Fig. 2C). Gene ontology (GO) analysis showed altered expression of genes involved in homeostasis of number of cells, RNA polymerase II transcriptional factor binding, and cardiac muscle cell differentiation/contraction (Supplementary Fig. 4D,E). Furthermore, a comparison study revealed that two-thirds of the downregulated genes in E10.5 hearts overlapped with an increase in H3K9me3 of the corresponding genes in *Cdk13* KD MEFs (Supplementary Fig. 4F). Immunofluorescence analysis showed a significant increase in known heterochromatin markers, such as H3K9me3, HP1β, HP1γ, and Trim28/KAP-1, in E13.5 *Cdk*13*^-/-^* embryonic hearts compared to control hearts (Fig. 2D-H). Similarly, changes in heterochromatin markers were observed in E10.5 *Cdk*13^-/-^ hearts (Supplementary Fig. 5A-D). In contrast, there was a decrease in phosphorylated RNA polymerase II at Ser2 and Ser5 was evident at E10.5, E11.5, and E13.5 hearts (Supplementary Fig. 6A-F). Additionally, we examined whether *Cdk*13^-/-^ embryonic heart defects were linked to changes in cell proliferation. The results showed a significant reduction in Ki67-positive cells in E9.5 *Cdk*13^-/-^ hearts (Supplementary Fig. 6G, H), compared to controls. Furthermore, the total number of cells in the E13.5 *Cdk*13^-/-^ mutant embryos was decreased by almost 50% (Supplementary Fig. 6I). These results support the critical role of Cdk13 in development and confirm Cdk13’s role in preventing heterochromatin expansion and specific gene repression *in vivo*.

## Phosphoproteomic analysis of the proteins caused by *Cdk*13 knockdown

To investigate how Cdk13 regulates the spreading of H3K9me3-heterochromatin, we hypothesized that Cdk13 may phosphorylate heterochromatin regulators, which in turn alters their interactions and subsequent heterochromatin spreading. We conducted phosphoproteomic analysis on *Cdk*13-KD MEFs and control MEFs (Fig. 3A). Data from replicates of cells transfected with *Cdk*13 shRNA showed 1673 downregulated phosphorylation sites within 976 proteins (Fig. 3B). Among the proteins with reduced phosphorylation, 69% are nuclear proteins, 15% are cytoplasmic proteins, and 3% are both nuclear and cytoplasmic proteins (Fig. 3C). Over 620 proteins carry one downregulated phosphorylation site, while the remaining proteins possess two or more downregulated sites (Fig. 3D), with highly conserved motif. The top three most matched preferential motifs, _xx__S_P_xx_, _xx_R_xx__S__xx_ and _xx__S__x_E_xx_, enriched from 1639 unique peptide sequence centered by downregulated phosphorylation site and extended 6 amino acid up- and downstream (Fig.3E). Compared with control, the changes in the phosphorylation of specific proteins in overexpression or depletion of *Cdk*13 were confirmed by Western blot (Supplementary Fig. 7B). The results support the significant role of Cdk13 in phosphorylation regulation, with a preference for targeting nuclear proteins.

**Fig. 3.**
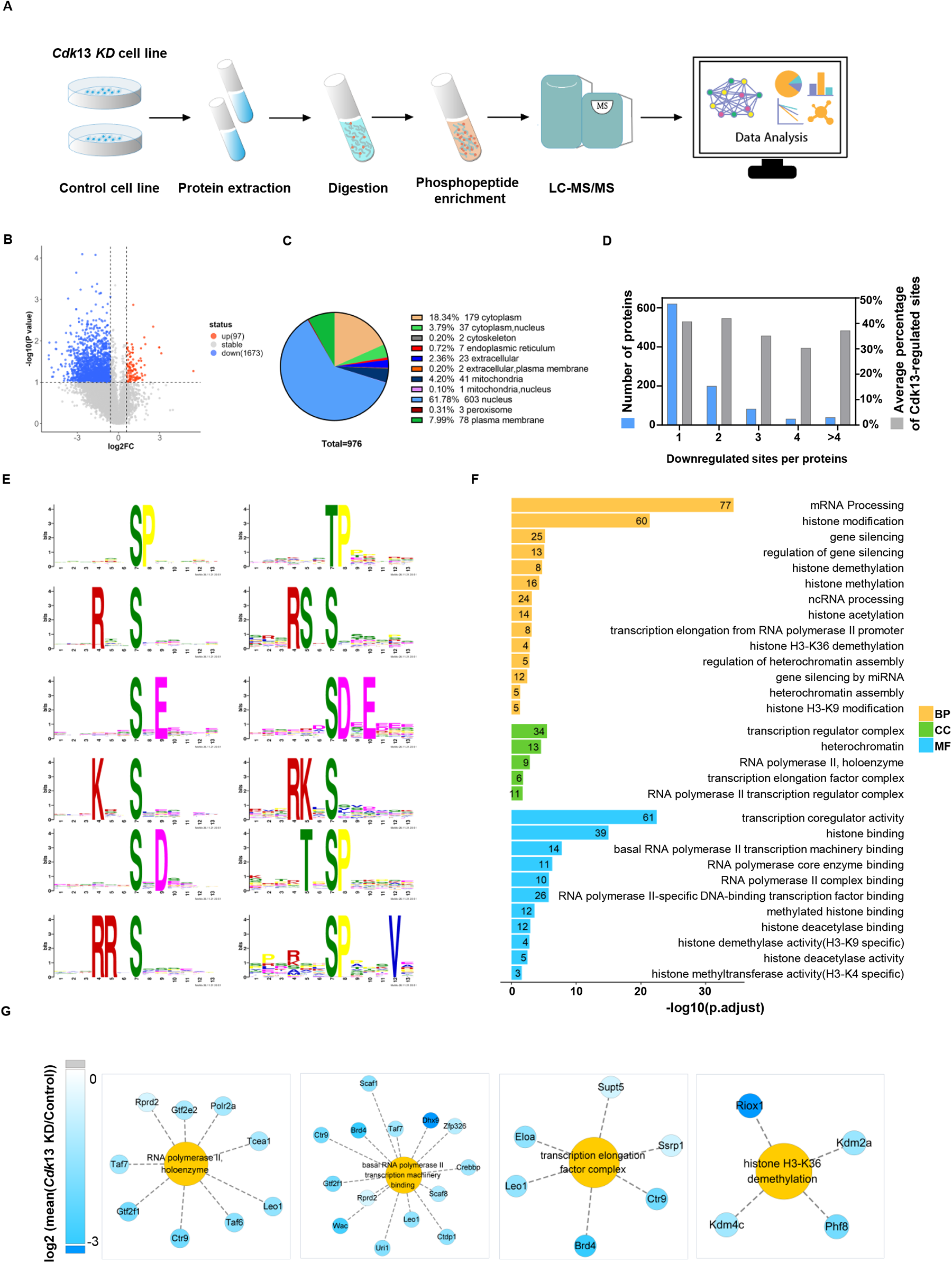
Phosphoproteomic analysis of the global protein network targeted by Cdk13. **A**, Global phosphoproteomics workflow. **B**, Majority changes in protein phosphorylation are downregulated. The volcano plot shows the detected S/T/Y sites in phosphorylation data. The X-axis represents log2(fold change), while the Y-axis represents -log10(P values). Red indicates upregulated sites, Blue indicates downregulated sites, and dotted lines represents the threshold of fold change (1.5) and P value (0.1). Each point represents a S/T/Y site. **C**, The pie plot shows the locations of downregulated proteins (fold change>1.5 and P < 0.1), most of these proteins enriched in the nucleus. **D**, More than 30% protein phosphorylation sites are downregulated in each group. The bar plot shows the relationship between the number of proteins (blue, left y-axis) with downregulated protein phosphorylation sites, the percentage of downregulated sites (grey, right y-axis), and the number of downregulated protein phosphorylation sites in each protein (x-axis). **E**, The amino acid sequence logo illustrates the sequence preference around downregulated phosphorylation sites after *Cdk*13 KD. The motifs were centered on the downregulation sites and extended 6 positions upstream and downstream from 1639 unique peptide sequences. **F**, Gene Ontology enrichment analysis of proteins with reduced phosphorylation after *Cdk*13 KD. Selected terms for each group in the BP, CC and MF categories are shown. Yellow represents biological process (BP); green represents cellular component (CC); blue represents molecular function (MF). The x-axis represents the -log10 (adjusted *P* value), and the numbers on the color bars indicate the numbers of genes enriched for each term. **G**, Proteins with loss of phosphorylation are associated with RNA polymerase II initiation and elongation, as well as regulators of specific histone modifications.

The top proteins found to have altered phosphorylation are involved in messenger RNA processing/splicing, and histone modifications (Fig. 3F). Specifically, more than 60 proteins possess transcription coregulation activity: 26 are RNAPII-specific DNA binding transcription factors, 14 are related to basal RNAPII transcription machinery binding, 9 are RNAPII holoenzymes, and 11 are related to RNAPII core enzyme binding (Fig. 3F,G). Many regulators of histone binding, histone methylation/demethylation or acetylation/deacetylation and gene silencing were identified (Supplementary Fig. 7A).

Furthermore, more than 20 of the identified proteins with downregulated phosphorylation are associated with heterochromatin assembly and H3K9 methylation or demethylation (Fig. 4A, Supplementary Fig.7A). For instance, Setdb1, an HMT that catalyses H3K9me3, and two regulators of Setdb1, Resf1 and ATFiP (Fig. 4A, B), were identified among the proteins with downregulated phosphorylation sites. Therefore, Cdk13 likely prevents heterochromatin spreading via phosphorylation on H3K9 HMTs. Moreover, the loss of phosphorylation at two specific sites of the heterochromatin protein HP1β, S89 and S91, was observed (Fig. 4B). Both sites are located within the Hinge region of HP1β, which is known to possess DNA/RNA binding activity. These results therefore support the notion that heterochromatin remodelers are targets of Cdk13, which may in turn contribute to the dynamics of heterochromatin spreading. Overall, the phosphoproteomic analysis shows that Cdk13 is a key regulator of proteins related to RNAPII transcription, most interestingly, Cdk13 was found to regulate the phosphorylation of drivers involved in heterochromatin formation.

**Fig. 4.**
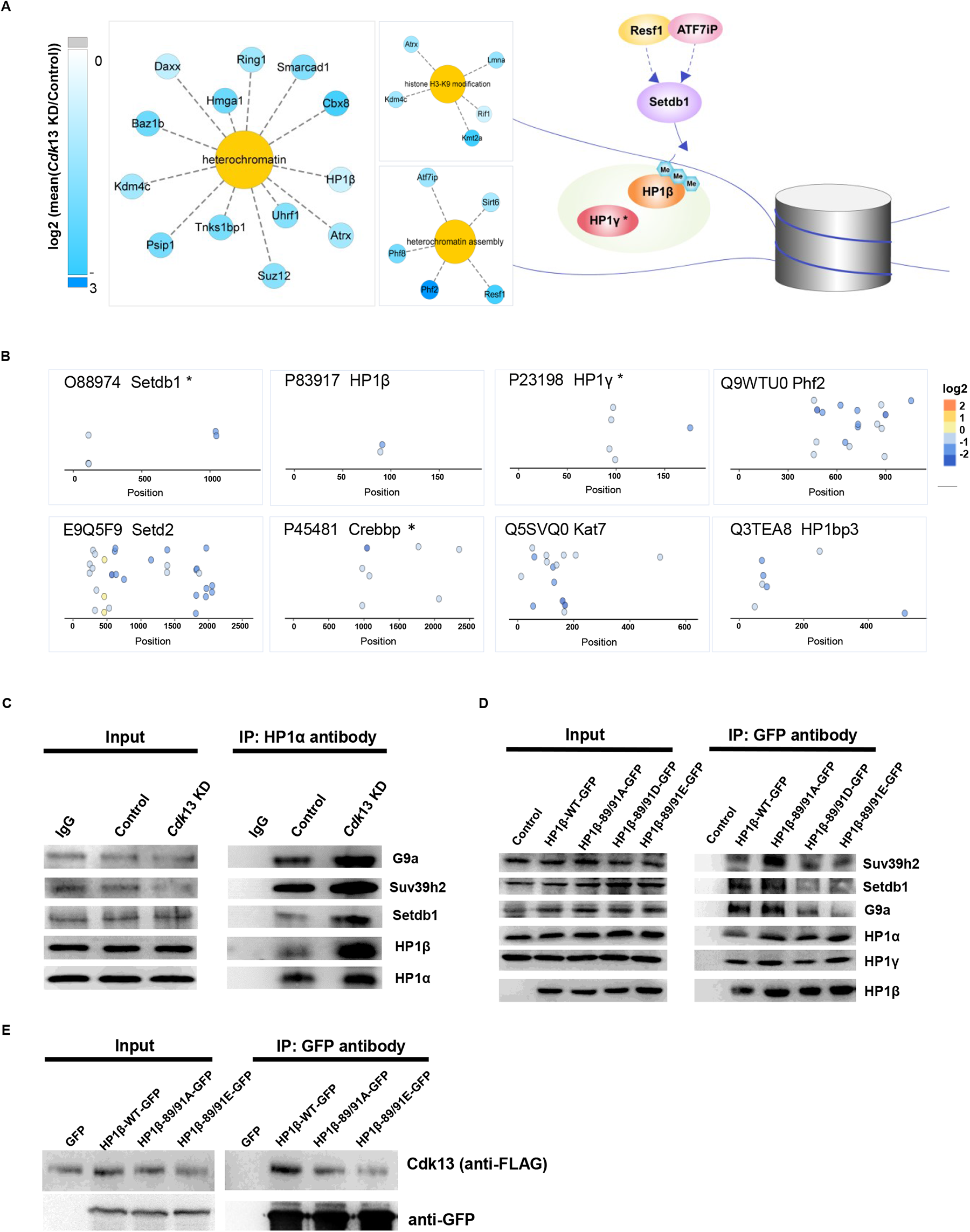
Inhibition of heterochromatin complex formation by specific phosphorylation of HP1β. **A**, Heterochromatin proteins with loss of phosphorylation and a possible model of heterochromatin assembly according to the results of global phosphoproteomics. **B**, The dot plot shows the change in representative protein phosphorylation sites; each point represents an amino acid (S/T) as a substrate for phosphorylation. The X-axis indicates the location of the phosphorylation site within the protein. The names of the selected proteins, including heterochromatin proteins H3K9me3 HMT-Setdb1 and HP1β, are indicated, and Log2 values indicate the site-specific phosphorylation ratio of *Cdk*13 KD/Control. **C**, Coimmunoprecipitation with an HP1α antibody followed by Western blot analysis was done to analyse the interactions of HP1α with G9a, Suv39h2, Setdb1, and HP1β in mouse embryonic fibroblasts (MEFs) infected with lentiviruses bearing shRNA in control or *Cdk*13 KD group. **D**, Phosphorylation at Ser89/91 of HP1β inhibited the binding of HP1β to the H3K9-associated methyltransferases, G9a, Suv39h2, and Setdb1. **E**, Co-immunoprecipitation revealed an interaction between Cdk13 and HP1 β, which was inhibited by HP1β phosphorylation. The anti-Flag antibody was used to monitor the presence of Cdk13, and the anti-GFP antibody was used to immunoprecipitate the GFP-tagged wild type or mutant of HP1β.

## Phosphorylation of HP1 attenuates the interaction between the master regulators of heterochromatin

To evaluated whether the affected phosphorylation of nuclear proteins participated in *Cdk*13 depletion-induced heterochromatinization and target gene suppression, we chose the heterochromatin protein HP1 for the study. HP1 recognizes H3K9me2/3 and can also serve as a critical ‘hub’ to recruit other heterochromatin modifiers and orchestrate the formation of heterochromatin (Gerwal and Jia 2007; Eissenberg and Elgin 2014). Firstly, we conducted immunoprecipitation experiments to examine the interaction between heterochromatin proteins such as HP1 orthologues, and the H3K9me2/3 HMTs Suv39h2, G9a, and Setdb1 after *Cdk*13 KD. Protein immunoprecipitation was performed using an antibody against HP1 orthologue and nuclear extracts from control and *Cdk*13-KD MEFs. The results indicated a significant increase in the affinity between HP1α and HP1β as well as H3K9me HMTs after *Cdk*13 depletion (Fig. 4C), supporting phosphorylation by Cdk13 prevents the heterodimerization.

We then examined whether the phosphorylation of specific sites on HP1, detected in the phosphoproteomic analysis, played a role in the interaction with H3K9me HMTs. To mimic phosphorylation loss or constitutive phosphorylation, we generated specific point mutations at S89 and S91 in HP1β (Fig. 4B,D), and green fluorescent protein (GFP) tags were fused to the C-terminus of wild-type HP1β and each mutant HP1β (Fig. 4D). Using the same immunoprecipitation analysis approach as described above, the results showed that the HP1β mutation carrying either 89/91D or 89/91E reduced the interaction between HP1β and the HMTs Suv39h2, Setdb1 and G9a (Fig. 4D). This suggests that constitutively phosphorylated HP1β loses its capability as a “hub” in orchestrating heterochromatin protein complexes, while the 89/91A mutation, mimicking the loss of phosphorylation, increases the interaction between HP1β and HMTs (Fig. 4D), which eventually promotes heterochromatin spreading. To support our hypothesis, we performed IP experiments using MEFs co-transfected with Flag-tagged Cdk13 and the HP1β mutant plasmid. The IP results showed an interaction occurring between the wildtype of Cdk13 and HP1β. However, the interaction between phosphorylated HP1β, as mimicked by mutation at S89 and S91 – HP1β-89/91EGFP, and Cdk13, was dramatically reduced (Fig. 4E).

## Inhibiting H3K9me3 prevents the expansion of heterochromatin and transcriptional gene silencing triggered by *Cdk*13 depletion

If the spreading of H3K9me3-heterochromatin was the cause of the observed defects in the *Cdk*13 mutant embryos, we hypothesized that reducing the level of H3K9me3 could potentially reverse the *Cdk*13 KD-induced HP1β association with chromatin and heterochromatin spreading. This could lead to the recovery of normal transcription levels and potentially relieve the developmental defects caused by the loss of *Cdk*13. Therefore, we conducted a pilot screen to identify an inhibitor of H3K9me3-marked heterochromatin spreading in the *Cdk*13 depletion background. From the chemical compound library, we selected fifteen inhibitors that target various activities, including histone H3K9me HMTs, repressors of active H3K4 methylation – LSD1, and HDAC inhibitors. After screening, we identified four inhibitors that significantly reduced H3K9me3 and increased the acetylation of H3K9ac (Fig. 5A, B; Supplementary Fig.8A, B). Among the four inhibitors, Givinostat (García-Rodríguez et al. 2020) (GIVI) is a pan-HDAC inhibitor of HDAC1 and HDAC3, while the other two are LSD1 inhibitors, and the remaining one is a G9a inhibitor (Supplementary Table 3). These findings are not surprising, as G9a is a known H3K9me2/3 HMT (Shinkai et al. 2011), and HDACs, along with LSD1, are components of repressive protein complexes (Shi et al. 2005) contributing to the crosstalk between histone modifications (Fischle et al. 2003; Ji et al. 2019). To further investigate these interactions, we conducted additional IP experiment and confirmed that HDAC3 interacts with HP1β complexes (Supplementary Fig. 8C), and the phosphorylation of HP1β blocked the interaction.

**Fig. 5.**
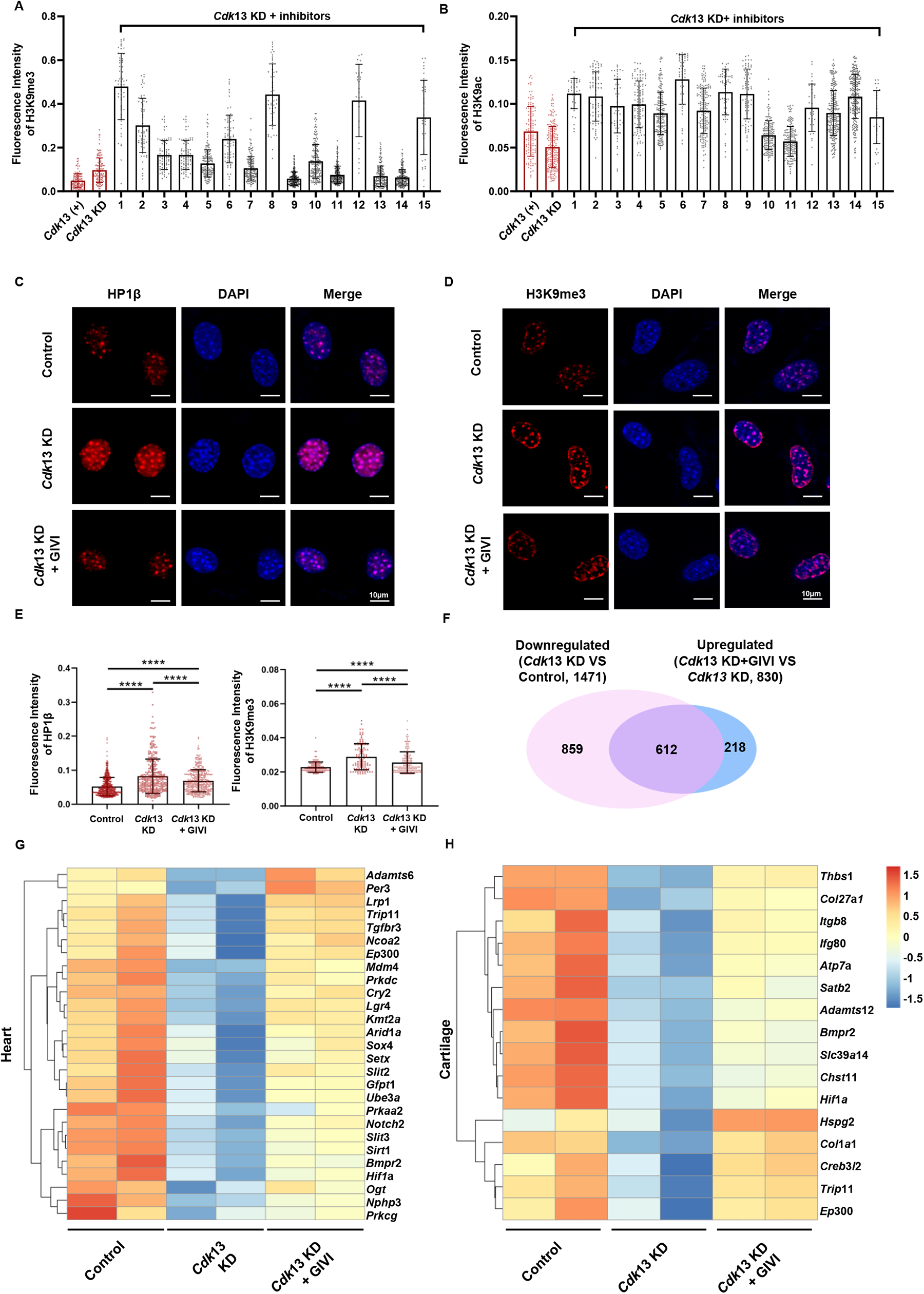
The depletion of *Cdk*13-induced heterochromatinization was antagonized by H3K9me3 inhibitors. **A, B** Fifteen small molecule inhibitors were screened by immunofluorescence staining assay to inhibit H3K9me3 and increase H3K9ac in *Cdk*13 KD cells. The left panel shows the quantification of fluorescence intensity of H3K9me3, and the right panel shows the quantification of H3K9ac fluorescence intensity. Each point represents a cell, and 30-250 cells per condition were analyzed. The data are presented as means ± SEMs. **C**, **D**, The small molecule drug givinostat (GIVI) (2.5 nM) blocked heterochromatin formation caused by *Cdk*13 knockdown. The representative immunofluorescence staining for HP1β (**C**) and H3K9me3 (**D**) in MEFs is shown. **E**, The quantification of the relative intensity of HP1β and H3K9me3 is shown. Each point represents a cell, and more than 300 cells per condition were analyzed. The data are presented as the means ± SEMs. \*\*\*\**P* < 0.0001, t-test. **F**, The GIVI treatment could efficiently recover more than 40% *Cdk*13 KD downregulated genes. The overlap between downregulated genes in *Cdk*13-depleted MEFs and upregulated genes after treatment with 2.5 nM GIVI is shown. **G**, **H**, RNA sequencing analysis of control, *Cdk*13 KD and *Cdk*13 KD +GIVI showed rescue trends. A heatmap shows several differentially expressed genes (DEGs) related to heart function (**G**) and cartilage (**H**).

The histone deacetylase inhibitor GIVI significantly inhibited *Cdk*13 KD-triggered H3K9me3 spreading, which also reduced the association of HP1β on chromatin (Fig. 5C-E, Supplementary Fig.8A) and elevated the active chromatin marker H3K4me3 (Supplementary Fig.9A,B). It is notable that the concentration of GIVI used, at 2.5nM, was 800-fold lower than that of LSD1 inhibitors and 400-fold lower than that of G9a inhibitors (Supplementary Table 3), yet it efficiently prevented H3K9me3-heterochromatin and recovered H3K9ac. Further RNA sequencing analysis showed that the inhibitor GIVI significantly recovered the transcription of approximately 42% of genes (612 in 1471) downregulated by *Cdk*13 depletion alone (Fig. 4F, Supplementary Fig. 9C). Most interestingly, among the rescued genes (Supplementary Fig. 9C), numerous targets were implicated in heart, cartilage and craniofacial development (Fig. 4G,H, Supplementary Fig. 9D).

## The parameters of heart defects caused by *Cdk*13 deletion are rescued by GIVI in mice

To investigate whether the reduction in H3K9me3 triggered by GIVI can affect the phenotype caused by the loss of *Cdk*13, we generated mice with heart-specific deletion of *Cdk*13 by crossing *Cdk*13*^fl/fl^* mice with myosin-heavy chain 6 (*mhc*)-cre mice to obtain *Mhc-cre;Cdk*13*^fl/fl^* mice. Unlike *Cdk*13*^-/-^*mice, which normally died around E14 with atrial defects (Fig. 2A), the *Mhc-cre;Cdk*13*^fl/fl^*mice were viable after birth, whereas lethality started to be observed at 159 days after birth, and all of these mice died before 260 days of postnatal life. In comparison, the median longevity of *Mhc-cre* control mice was 329 days under the same animal feeding conditions. Histological analysis was carried out to examine whether lethality was caused by postnatal heart defects. Indeed, the results showed a thin atrioventricular wall, heart enlargement and atrial congestion caused by dilation of the ventricles and atria (Fig. 6A), and a fourfold increase in myocardial fibrosis in the *Mhc-cre;Cdk*13*^fl/fl^*mice at 8 months of age compared with controls (Supplementary Fig. 10A). At this stage, the *Mhc-cre;Cdk*13*^fl/fl^* mice also showed significantly decreased left ventricular posterior wall (LVPW) thicknesses at diastole (d) and systole (s), increased left ventricular internal dimensions (LVID) at diastole (d) and systole (s) (Supplementary Fig. 10B, upper panel), and an enlarged left ventricular volume (Supplementary Fig. 10B, lower panel). In addition, left ventricular global systolic function was significantly reduced, as demonstrated by the reduced left ventricular ejection fraction (EF) and left ventricular fractional shortening (FS) (Supplementary Fig. 10B, lower panel), supporting the requirement of Cdk13 in the heart structure and function.

**Fig. 6.**
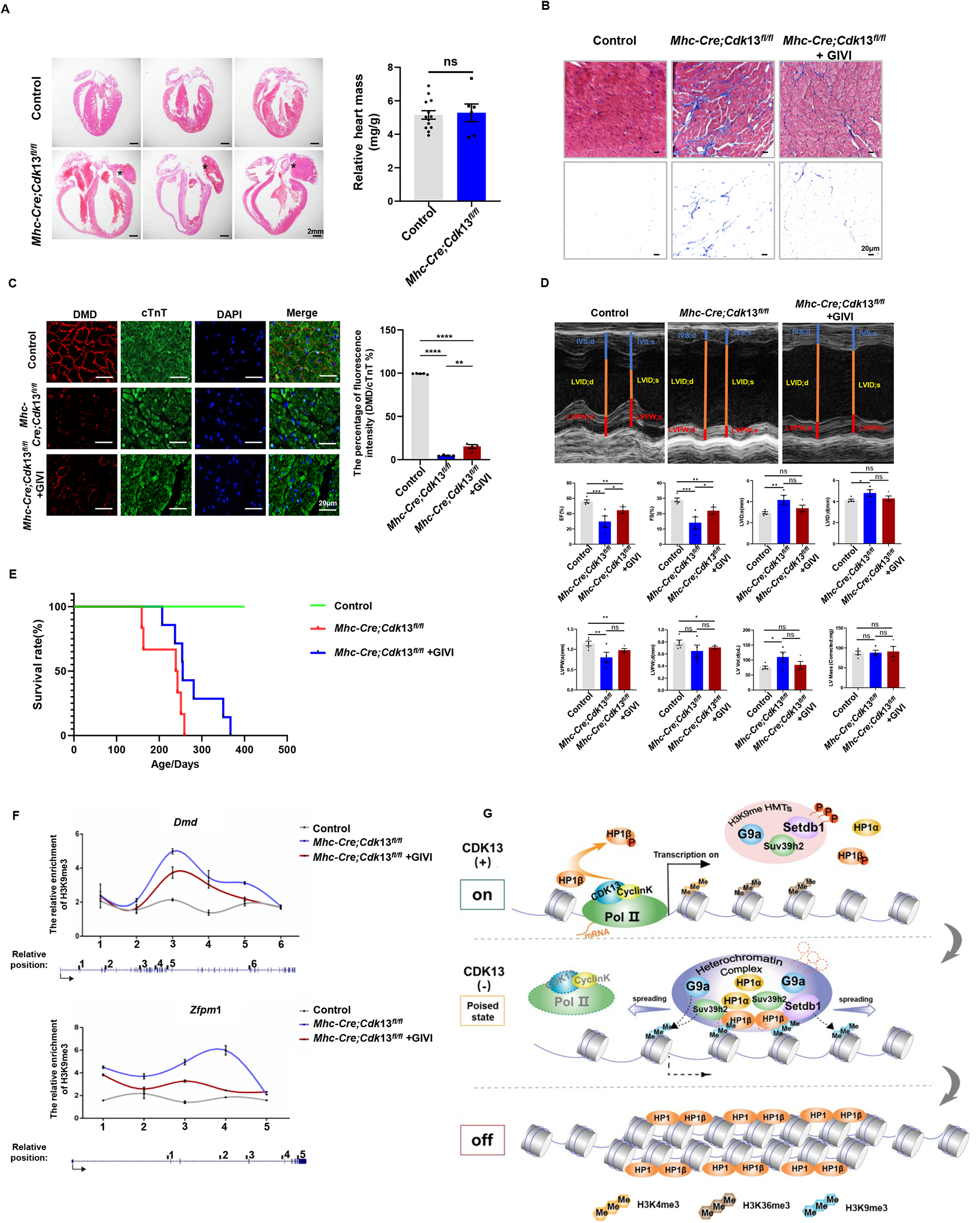
Heart defects caused by *Cdk*13 deletion in mice are alleviated by GIVI. **A**, Ventricular lumen enlargement and ventricular wall thinning were observed in 8-month-old *Mhc-cre;Cdk*13*^fl/fl^*mice. Haematoxylin and eosin were used to illustrate the heart structure. Asterisks indicate atrial congestion. The right panel shows the statistics of the heart weight (mg) relative to body weight (g). **B**, GIVI partially reversed fibrosis caused by *Cdk*13 knockout. The hearts of mice stained with Masson’s stain to indicate the pathological fibrosis process. The scale bar was 50 μm. **C**, GIVI or DMSO treated heart tissues were stained with antibodies against Dmd (red) and cTnT (green). The right panel shows the percentage Dmd and TnT-positive cells by fluorescence intensity analysis, *n* = 4. \*\**P* <0.01,\*\*\*\**P* < 0.0001, t-test. **D**, echocardiographic tracing of mouse hearts. IVS indicates interventricular septum. LVID indicates left ventricular internal diameter. LVPW indicates left ventricular posterior wall. EF indicates ejection fraction; FS indicates fractional shortening; LVPW indicates left ventricular posterior wall, systolic (s); LVPW indicates left ventricular posterior wall, diastolic(d); LV mass indicates left ventricular mass. **E**, The survival rate of GIVI-treated *Mhc-cre*;*Cdk*13*^fl/fl^*mice was significantly increased compared to the *Mhc-cre*;*Cdk1*3*^fl/fl^*group without GIVI treatment. **F**, ChIP-qPCR analysis showed that the enrichment of H3K9me3 in *Cdk*13 knockout embryos was decreased in the presence of GIVI. The E17.5 heart tissue of control (*Cdk*13*^fl/fl^*), *Cdk*13*-*deletion mutants (*Mhc-cre;Cdk*13*^fl/fl^*), and the GIVI-treated *Mhc-cre*;*Cdk*13*^fl/fl^*mice were collected to perform the ChIP experiment, followed by qPCR analysis. The primers were designed to amplify the defined regions in the *Dmd*, and *Zfpm*1 genes. **G**, Graphic model of Cdk13 in gene transcription and heterochromatin formation. The Top panel shows Cdk13 maintains normal transcription activation by phosphorylating heterochromatin proteins. The middle panel shows that the deletion of *Cdk*13 causes a loss of HP1β phosphorylation, which increases the interaction between HP1β and heterochromatin proteins, leading to the spreading of poised heterochromatin structure at the euchromatic gene region. At the bottom, a stable repressive facultative heterochromatin structure is formed, resulting in the subsequent gene silencing. ns, not significant: P > 0.05; *P < 0.05; **P < 0.01;***P < 0.001; ****P < 0.0001, t-test.

The defect in myocardial function was accompanied by downregulation of numerous genes associated with striated muscle differentiation, striated muscle contraction, contractile fibres, regulation of heart contraction force, cardiac muscle cell action potential, and regulation of blood circulation (Supplementary Fig. 10C-F), in the left ventricle and left atrium in 3.5-month-old *Mhc-cre;Cdk*13*^fl/fl^* mice. Several important genes associated with myocardial development, such as *Bmp*7, *Zfpm*1, *Hand*2, *Tbx*6, and *Dmd,* were found among the downregulated genes (Supplementary Fig. 10E, F). Specifically, Bmp7 contributes to ventral body wall closure (Nishimatsu and Thomsen 1998; Schmid et al. 2000; Kim et al. 2019), Zfpm1 is involved in ventricular septal defects (Katz et al. 2003), Hand2 is required for cardiac gene expression and outflow tract development (Holler et al. 2010; VanDusen et al. 2014), and Tbx6 is critical for cardiovascular lineage diversification (Sadahiro et al. 2018). Mutation of *Dmd* is a well-known genetic disease caused by the loss of functional dystrophin protein, which is responsible for the stability of skeletal muscle fibres and cardiomyocytes during contraction (García-Rodríguez et al. 2020; Mercuri et al. 2013). The transcriptional reduction of *Dmd* was also confirmed by the immunofluorescence staining in left ventricular tissues from 7-month-old *Cdk*13*^fl/fl^*mice (Supplementary Fig. 10G). The reductions in the expression of *Dmd* and other genes in *Cdk*13*^fl/fl^* mice perfectly illustrated the myocardial dysfunction.

To study whether the heart defects and epigenetic gene silencing in *mhc-cre;Cdk*13*^fl/fl^* mice were the consequence of *Cdk*13 mutation in prenatal development, *Cdk*13 was conditionally deleted in postnatal mice generated by crossing *Cdk*13*^fl/fl^*mice with *ubc-creERT2* mice. The offspring *ubc-creERT2;Cdk*13*^fl/fl^*mice, which expressed cre recombinase in most tissues after tamoxifen induction for 5 days at 1 month of age, were designated *Cdk*13*^uKO^*mice. After 12 months of postnatal life, the *Cdk*13*^uKO^* mice developed normally, with no apparent differences in heart morphology, behaviour or body weight compared with those of controls (Supplementary Fig. 10H). The deletion efficiency of *Cdk*13 in *creERT2*;*Cdk*13*^fl/fl^*mice was measured using standard PCR, and approximately 90% deletion was observed (Supplementary Fig. 10I). This result, along with the early embryonic heart defects caused by the loss of *Cdk*13, supports that Cdk13 is a critical regulator of prenatal development.

We next evaluated whether inhibition of H3K9me3 by GIVI can rescue the mouse heart defects caused by the loss of Cdk13. *Mhc-cre;Cdk*13*^fl/fl^*mice at 6–7 months of age were treated with 10 mg/kg GIVI per day for 30 consecutive days. Immunohistological analysis of the left ventricle showed a reduction in the fibrotic area in GIVI-treated *Mhc-cre;Cdk*13*^fl/fl^* mice (Fig. 6B). In addition, the expression of *Dmd* was also elevated after GIVI treatment (Fig. 6C). Further echocardiogram analysis showed that markers of left ventricle global systolic function in *Mhc-cre;Cdk*13*^fl/fl^* mice, such as the left ventricular ejection fraction and left ventricular fractional shortening, were significantly recovered by GIVI treatment (Fig. 6D). Notably, the survival rate of GIVI-treated *Mhc-cre*;*Cdk*13*^fl/fl^* mice was significantly increased compared to the *Mhc-cre*;*Cdk*13*^fl/fl^*group without GIVI treatment, with a maximum lifespan extended more than 40% (108 days) (Fig. 6E). These results suggest that inhibiting H3K9me3 by GIVI is sufficient to alleviate heart defects caused by the loss of Cdk13 and facilitates the survival rate of mutant animals.

To confirm any changes in H3K9me3 on chromatin in GIVI-treated *Mhc-cre*;*Cdk13^fl/fl^*mice, we performed ChIP quantitative PCR to examine the enrichment of H3K9me3 on the *Dmd* gene and two other known key regulators of myocardial development, *Bmp*7 and *Zfpm*1. We used heart tissue from *Mhc-cre*;*Cdk*13*^fl/fl^*, GIVI-treated *Mhc-cre*;*Cdk*13*^fl/fl^*, and control mice at E17.5 for the ChIP quantitative PCR. The results showed that the elevations in H3K9me3 intensity along the majority of gene bodies of *Dmd*, *Bmp*7, and *Zfpm*1 in *Mhc-cre*;*Cdk*13*^fl/fl^*mice were indeed reduced in the presence of GIVI (Fig. 6F, Supplementary Fig. 10J), further supporting the key role of H3K9me3 in *Cdk*13-congenital syndrome and suggesting that GIVI is a potential drug for treating the genetic disease.

## Discussion

There are increasing studies indicating the important role that metazoan C-terminal domain (CTD) kinase members – dCDK12/Cdk12/hCDK12 and Cdk13/hCDK13 – play in transcription and diseases. In the genome of fruit flies – *Drosophila melanogaster* – the orthologue is named dCDK12 (Bartkowiak et al. 2010). In humans, it is known as CDK12/hCDK12 and CDK13/hCDK13 (Chen et al. 2006). Several studies have reported that dCDK12/CDK13/hCDK13 is involved in CTD phosphorylation at Ser2/5 (Pan et al. 2015; Bartkowiak et al. 2010; Greifenberg et al. 2016), RNA polymerase II processivity or nuclear RNA surveillance (Fan Z et al., Sci Adv. 2020; Insco ML et al., 2023, Science).

We demonstrated in this study that mice Cdk13 plays an important role in preventing heterochromatin spreading. We further show that Cdk13 modulates the phosphorylation of master heterochromatin regulators, including H3K9me HMTs – Suv39h2, Setdb1, and G9a – and the reader of H3K9me2/3 – HP1 orthologues. The loss of specific HP1β phosphorylation at S89 and S91 increases the interaction between HMTs and HP1β (Fig. 4C, D). As a result, this may lead to heterochromatin spreading at euchromatic genes, both in *Cdk*13-KD MEFs and *Cdk*13*^-/-^* mutant tissues, and various embryonic defects (Fig. 1, Fig. 2D-G, Fig. 6F, Supplementary Fig. 3). The observation that Cdk13 mainly affected the development of prenatal mice (Supplementary Fig.3C-F), but appeared to be redundant in postnatal mice (Supplementary Fig.10H),suggests a temporospatial role of mice Cdk13. This is also supported by the fact that approximately 90% deletion was observed in adult *creERT2*;*Cdk*13*^fl/fl^* mutant animals (Supplementary Fig. 10I), and other phenotypic analysis with different *cre*-drivers (data not shown).

In conclusion, our data reveal that Cdk13 plays a critical role in a phosphorylation-based mechanism for facultative H3K9me3-heterochromatin spreading in mammals. The present study also suggested that H3K9me3-heterochromatin spreading is a key driver of *Cdk*13 mutation-induced congenital heart syndrome. The identified GIVI has been found to relieve congenital heart defects triggered by *Cdk*13 mutation, which may provide a theraputic approach for *CDK*13 mutation or heterochromatin alteration-induced congenital diseases.

## Methods

### Mouse usage, Embryonic development, and Morphology identification

All experiments and maintenance of mice in this study were conducted in compliance with the University of Health Guide for the Care and Use of Laboratory Animals and approved by the Biological Research Ethics Committee of Tongji University.

For mice with different gestation periods, female mice were euthanized by cervical dislocation, and their abdomens were disinfected with 70% ethanol. The uterus, full of embryos, was then removed, and individual embryos were separated from the uterine membrane. The embryos were observed and photographed under a stereomicroscope. Observation of the vaginal plug was considered Embryonic Day 0.5 (E0.5). The genetic background of wildtype and the *Cdk*13 mutant genetic background of Cdk13^fl/fl^ was C57/BL6.

### Cell culture

MEF/3T3 cells were cultured in Dulbecco’s Modified Eagle’s Medium (catalog number R10-017-CV) from Corning, NY, USA, containing 10% fetal bovine serum (catalog number 16000) from Gibco, Waltham, MA, USA, and 1% penicillin/streptomycin (catalog number SV30010) from HyClone, Logan, UT, USA. The cells were cultured in a 37 °C incubator with 5% CO2.

### shRNA

The targeting sequences against the Cdk13 gene used in the present study were as follows: Cdk13 KD (5’-AGAGTATCATCAATATGAA-3’) and control (5’-TTCTCCGAACGTGTCACGT-3’). They were cloned into the hU6-MCS-Ubiquitin-EGFP-IRES-puromycin vector and packaged into lentiviruses. The packaged lentiviruses harboring the shRNA vectors were then introduced into mouse embryonic fibroblasts, and flow cytometry was used to separate the GFP-positive cells to obtain stable cell lines.

### Immunocytochemical staining

Mouse embryonic fibroblast cells were grown on slides until they were adherent. The cells were fixed with 4% paraformaldehyde (w/vol) for 30 min at 25 °C, and washed with phosphate-buffered saline (PBS) three times, 10 min for each. Then cells were permeabilized with 0.1% Triton-X 100 in PBS for 30 min at 25 °C, washed with PBS three times, and then blocked with a solution containing 4% bovine serum albumin for 1 h at 25 °C. Primary antibodies against HP1β (ab10478) (Abcam, Cambridge, UK), H3K9me3 (ab8898) (Abcam, Cambridge, UK), H3K4me3 (ab8580) (Abcam, Cambridge, UK), H3K9ac (ab10812, Cambridge, UK), H3K27ac (ab4729, Cambridge, UK), Trim28 (ab109287, Cambridge, UK) and H4K16ac (cs204361, Millipore, USA) were added, and the cells were incubated at 4 °C overnight. The cells were then washed with PBS three times.

A secondary antibody solution was dropped onto the cells, and they were incubated for 1 hour at 25 °C in the dark. Then, the cells were washed with PBS three times. DAPI (5 μg/ml) was added, and the cells were incubated at room temperature in the dark for 10 minutes. The cells were then washed three times in PBS. Finally, anti-fluorescence quenching agent was added to the slides to seal them, and fluorescence photographs were taken.

### Transmission electron microscopy

*Cdk*13 KD and control mouse embryonic fibroblast cells were digested with 0.125% trypsin. The digestion was blocked with serum, and the cells were centrifuged at 350 g for 5 min. The cell pellets were fixed with 2.5% glutaraldehyde in PBS (0.1 M phosphate buffer, pH7.4) for 30 min at 25°C. The cell precipitates were embedded and sectioned following standard procedure, and were examined under transmission electron microscopy.

### Preparation of tissue into paraffin sections

Freshly removed tissues were washed with PBS to remove blood, the 4% paraformaldehyde equal to the 20-fold volume that of tissue was added, and the tissues were fixed at 4 °C overnight. After washing with PBS three times, 70% ethanol was added, and the tissues were allowed to dehydrate overnight. The tissues were placed in an embedding box and dehydrated in a gradient of 80% ethanol, 90% ethanol, 95% ethanol I, 95% ethanol II, 100% ethanol I and 100% ethanol II for 30 min per concentration. The embedding box was placed in xylene II and xylene I for 3 min each. The embedding box was then placed in benzene wax, wax I, wax II and wax III for 30 min in sequence at 60 °C. After embedding, the tissue blocks were placed on a -20 °C cold table for at least 1 h. After solidification, paraffin sections with a thickness of 5 μm were prepared using a slicer. After spreading on a 42 °C slide machine, the sections were placed onto slides and incubated in a 37 °C oven overnight.

### Immunofluorescence staining of paraffin sections

The paraffin sections were placed in xylene I and xylene II for 5 min, and sequentially placed in 100% ethanol I, 100% ethanol II, 95% ethanol I, 95% ethanol II, 90% ethanol, 80% ethanol, 70% ethanol, ddH_2_O and PBS for 2 min each. The slides were placed in preheated sodium citrate (pH = 6.0) for 20 min at 97 °C. The slides were washed with PBS after the temperature dropped to 25 °C. Bovine serum albumin was dropped onto the slides, which were then left to stand at room temperature for 1 h. Primary antibodies against HP1β (ab10478) (Abcam, Cambridge, UK), HP1γ (ABE1329) (Millipore, Burlington, MA, USA), H3K9me3 (ab8898) (Abcam, Cambridge, UK), Trim28 (ab109287) (Abcam, Cambridge, UK), Ki67 (ab16667, Cambridge, UK), RNA polymerase II p-Ser2 (ab5095, Cambridge, UK), RNA polymerase II p-Ser5 (ab5131, Cambridge, UK) Dmd (ab15277) (Abcam, Cambridge, UK), and cTnt (MA5-12960) (Invitrogen, Waltham, MA, USA) were added, and the slides were incubated overnight at 4 °C before washing three times by PBS. The secondary antibody solution was dropped onto the slides, which were incubated for 1 h at room temperature in the dark and then washed with PBS three times. DAPI (5 μg/ml) was added onto the slides and incubated at room temperature under dark for 10 min, then washed in PBS three times. Anti-fluorescence quenching agent was dropped onto the slides to seal them, and then fluorescence photographs were taken.

### Haematoxylin and eosin staining

The paraffin sections were placed in xylene I and xylene II for 5 min and then sequentially incubated in 100% ethanol I, 100% ethanol II, 95% ethanol I, 95% ethanol II, 90% ethanol, 80% ethanol, 70% ethanol, and ddH_2_O for 2 min each. The sections were then washed with ddH_2_O three times. The sections were soaked in haematoxylin staining solution for 30 s, rinsed with ddH_2_O, soaked in 1% acetic acid for 30 s, rinsed with ddH_2_O, soaked in eosin staining solution for 2 min, and then rinsed with ddH_2_O. The sections were placed in 95% ethanol III, 95% ethanol IV, 100% ethanol III and 100% ethanol IV for 2 min each in sequence for dehydration. The sections were placed in xylene III and xylene IV for 5 min each in sequence, sealed with neutral gum and left to dry.

### Masson staining

The paraffin sections were sequentially placed in xylene I and xylene II, each for 5 minutes; 100% ethanol I and 100% ethanol II, 95% ethanol I and 95% ethanol II, 90% ethanol, 80% ethanol, 70% ethanol, and ddH_2_O for 2 minutes each, and then washed with ddH_2_O three times. The sections were soaked in Bouin’s staining solution and washed with ddH_2_O after incubation in a 56 °C bath for 15 minutes. Then, the sections were soaked in haematoxylin staining solution for 1 minute, rinsed with ddH_2_O, soaked in eosin staining solution for 30 seconds, and rinsed with ddH_2_O. The sections were then incubated in phosphomolybdic acid separation solution for 8 minutes and in aniline blue staining solution for 5 minutes before being rinsed with 1% acetic acid in ddH_2_O.

To dehydrate the sections, they were placed in 95% ethanol III and 95% ethanol IV, 100% ethanol III, and 100% ethanol IV for 2 minutes each. They were then placed in xylene III and xylene IV for 5 minutes each, sealed with neutral gum, and left to dry.

### Coimmunoprecipitation analysis

Mouse embryonic fibroblasts were collected and washed with ice-cold PBS, resuspended in RIPA lysis buffer (containing 25 mM Tris-Cl, pH 7.5; 150 mM NaCl; 1% NP-40; 1% sodium deoxycholate; 0.1% SDS; 1x protein cocktail; and 1 mM PMSF) and sonicated on ice for 4 min by 12% efficiency. The cell lysates were incubated with protein A/G beads (sc-2003) (Santa Cruz Biotechnology, Dallas, TX, USA) for 30 min to eliminate the background signal, and then incubated with anti-HP1α (NB110-40623) (Novus, St Louis, MO, USA) or GFP-Trap (gta-20) (Chromotek, Germany) antibodies at 4 °C overnight. The next day, for anti-HP1α binding, the samples were incubated in protein A/G agarose beads solution for 2 h. Then, the samples were centrifuged to harvest the agarose beads, and the beads were washed three times with immunoprecipitation buffer (20 mM HEPES, pH 8.0; 0.2 mM EDTA; 5% glycerol; 150 mM NaCl; and 1% NP-40). The precipitated proteins were released by boiling in loading buffer and separated by SDS–PAGE (10%). Immunoblot analyses were performed with antibodies against G9a (3306) (CST, Walall, UK), SUV39H2 (ab240313) (Abcam, Cambridge, UK), SETDB1 (A6145) (Abclonal, Woburn, MA, USA), HP1α (NB110-40623) (Novus, St Louis, MO, USA), HP1β (MAC-053) (Thermo Fisher, Waltham, MA, USA), HDAC3 (A16462, Woburn, MA, USA) and HP1γ (MAC-054) (Thermo Fisher, Waltham, MA, USA).

### RNA sequencing

Two micrograms of RNA were obtained and delivered to Novogene Company for RNA quality inspection, library construction and sequencing. The library type was a eukaryotic common transcriptome library (250-300 bp), the sequencing machine model was a HiSeq X Ten, the mode was PE150, and 9 GB of original data were obtained for each sample.

The sequencing files were mapped to the mouse genome mm10 using HISAT2 (v2.1.0) (Kim et al. 2015), and featureCounts (v1.6.2) (Liao et al. 2014) was used to obtain a gene expression matrix from the mapped reads. DESeq2 (v1.30.0) (Love et al. 2014) was used to identify differentially expressed genes (DEGs) with the LRT test. DEGs (differentially expressed genes) were defined as genes with an absolute fold change greater than 1.3 and an adjusted p-value less than 0.1. ClusterProfiler (v3.18.0) (Yu et al. 2012) and org.Mm.eg.db (v3.12.0) (Carlson et al. 2010) were used to perform Gene Ontology enrichment analysis.

### Chromatin immunoprecipitation assay

Mouse embryonic fibroblasts or mouse embryonic heart tissues were digested into single cells and washed with ice-cold PBS. The cells were incubated with 1% formaldehyde in PBS for 10 min, quenched with 0.125 M glycine, and washed twice with ice-cold PBS. The cell pellets were resuspended in ice-cold lysis buffer (50 mM Tris-Cl, pH 8.0; 140 mM NaCl; 1 mM EDTA; 10% glycerol; 0.5% NP-40; 0.25% Triton X-100; 1x protein cocktail), incubated on ice for 20 min, transferred to a prechilled Dounce homogenizer and homogenized. The nuclei were harvested by centrifugation, resuspended in nuclear lysis buffer (50 mM Tris-Cl, pH 8.0; 10 mM EDTA; 1% SDS; 1x protein cocktail) and incubated on ice for 20 min. The samples were sonicated on high power with 30s on and 30s off for a total 60 min. The solution of sheared chromatin was harvested and diluted with dilution buffer (50 mM Tris-Cl, pH 7.4; 150 mM NaCl; 1 mM EDTA; 1% NP-40, 0.25% deoxycholic acid). Then, 10 μg of anti-H3K9me3 antibody (ab8898) (Abcam, Cambridge, UK) was added, and the samples were incubated on a rotator overnight at 4 °C. Magnetic protein G beads (9006) (CST Pharma, Walsall, UK) were added, and the solution was incubated for 4 h. Then, the beads were harvested with a magnet and washed three times with wash buffer (100 mM Tris-Cl, pH 8.0; 500 mM LiCl; 1% NP-40; 1% deoxycholic acid; 1x protein cocktail). DNA was eluted with elution buffer (50 mM NaHCO_3_, 1% SDS) and extracted with a DNA purification kit. The eluted DNA was subjected to high-throughput sequencing or quantitative polymerase chain reaction.

For sequencing, DNA library construction and sequencing were performed by GENEWIZ Company. The sequencing files were mapped to the mouse genome mm10 using Bowtie2 (v2.4.2) (Langmead and Salzberg 2012) with the parameters --no-unal --no-mixed --no-discordant. The unmapped reads were filtered with samtools (v1.3.1) (Li et al. 2009) view with the parameters -F 12 -q 10, and duplicated reads were removed with sambamba (v0.8.1) (Tarasov et al. 2015) markdup. Mitochondrial DNA and blacklist regions were also removed. Macs2 (v2.2.7.1) (Zhang et al. 2008) was used for peak calling with the parameters -g mm -q 0.05 --broad -f BAMPE. The BAM files were converted into bigwig files using bamCoverage from deepTools (v3.5.1) (Ramirez et al. 2016) with the parameter -bs 10 --normalizeUsing RPKM. Signals around the TSS (±3 kb) were calculated and visualized with computeMatrix. The bigwig files were compared using bigwigCompare with -bs 10. The packages ChIPseeker (v1.22.1) (Yu et al. 2015) and TxDb.Mmusculus. UCSC.mm10.knownGene (v3.10.0) was used to annotate peaks. The protocols for Gene Ontology enrichment analysis were similar to those for RNA sequencing analysis.

Quantitative reverse transcription-polymerase chain reaction (qRT‒PCR) of H3K9me3 ChIP DNA was run in a Roche Light Cycler 96 thermocycler as follows: one cycle of 95 °C for 1 min; 40 cycles of 95 °C for 1 min, 60 °C for 1 min, and 72 °C for 1 min; and one cycle of 72 °C for 5 min. The data from the immunoprecipitates were normalized with the data from the 1% input samples. Please see the supplementary material for the primer sequence

### Echocardiography in mice

Cardiac function was evaluated using a high-resolution small-animal ultrasound imaging system (Vevo2100). Mice were anaesthetized with 1% isoflurane until the heart rate stabilized at 400– 500 beats per min. Seven parasternal long-axis images were obtained in B mode and scanned appropriately to determine the maximum left ventricular length. In this view, the M-mode cursor was perpendicular to the maximum left ventricular size at the end diastolic and systolic stages, and M-mode images were obtained to measure the wall thickness and chamber size.

### Treatment of cells and mice with small molecule drugs

For the cell experiment, the small molecule drugs were dissolved in DMSO to different concentration as a stock solution. The stock solution was added to *Cdk*13 KD mouse 3T3 cell growth medium in a final concentration (See Supplementary Table 3), and the cells were incubated for 72 h, then immunofluorescence experiments were performed.

For animal experiments, GIVI was dissolved in DMSO to 20 mg/ml and prepared as a stock solution. The 1 mg/ml working solution containing of 5% GIVI (stock solution), 40% PEG300, 5% Tween 80, and 50% ddH_2_O. The control solution containing of 5% DMSO, 40% PEG300, 5% Tween 80 and 50% ddH_2_O. Then, mice were given intraperitoneal injections of control and GIVI (10 mg/kg/day) solutions for 30 days.

### Protein phosphorylation mass spectrometry

Mouse embryonic fibroblasts, including *Cdk*13 knockdown (*Cdk*13 KD) and control cells (Control), were harvested (109 cells for each sample) and washed twice with PBS. Then, the samples were sonicated on ice for three minutes using a high-intensity ultrasonic processor (Scientz) in lysis buffer (8 M urea, 1% protease inhibitor cocktail). The remaining debris was removed by centrifugation at 12,000 g at 4°C for ten minutes. Finally, the supernatant was collected, and the protein concentration was determined using a BCA kit according to the manufacturer’s instructions. The sample was slowly added to the final concentration of 20% (m/v) TCA to precipitate protein, then mixed by vertexing and incubated for two hours at 4°C. The precipitate was collected by centrifugation at 4500 g for five minutes at 4°C. The precipitated protein was washed with pre-cooled acetone three times and dried for one minute. The protein sample was then redissolved in 200 mM TEAB and ultrasonically dispersed. Trypsin was added at a 1:50 trypsin-to-protein mass ratio for the first digestion overnight. The sample was reduced with 5 mM dithiothreitol for 60 minutes at 37°C and alkylated with 11 mM iodoacetamide for 45 minutes at room temperature in the darkness. Finally, the peptides were desalted using a Strata X SPE column.

Peptide mixtures were first incubated with IMAC microsphere suspension and vibrated in loading buffer (50% acetonitrile/0.5% acetic acid). To remove the non-specifically adsorbed peptides, the IMAC microspheres were washed sequentially with 50% acetonitrile/0.5% acetic acid and 30% acetonitrile/0.1% trifluoroacetic acid. To elute the enriched phosphopeptides, the elution buffer containing 10% NH_4_OH was added, and the enriched phosphopeptides were eluted with vibration. The supernatant containing phosphopeptides was collected and lyophilized for LC-MS/MS analysis.

The tryptic peptides were dissolved in solvent A (0.1% formic acid, 2% acetonitrile/ in water) and directly loaded onto a homemade reversed-phase analytical column (25 cm length, 100 μm i.d.). Peptides were separated with a gradient from 3% to 20% solvent B (0.1% formic acid in 90% acetonitrile) over 70 minutes, 20% to 30% in 12 minutes, and climbing to 80% in four minutes, then holding at 80% for the last four minutes, all at a constant flow rate of 500 nL/min on an EASY-nLC 1200 UPLC system (Thermo Fisher Scientific).

The separated peptides were analyzed in an Orbitrap ExplorisTM 480 (Thermo Fisher Scientific) with a nano-electrospray ion source. The electrospray voltage applied was 2.3 kV. The full MS scan resolution was set to 60,000 for a scan range of 400 -1200 m/z. Up to 15 most abundant precursors were then selected for further MS/MS analyses with 30 s dynamic exclusion. The HCD fragmentation was performed at a normalized collision energy (NCE) of 27%. The fragments were detected in the Orbitrap at a resolution of 30,000. Fixed first mass was set as 110 m/z. Automatic gain control (AGC) target was set at 75% with an intensity threshold of 2E4 and a maximum injection time of 100 ms.

### Protein phosphorylation data analysis

The resulting MS/MS data were processed using the Proteome Discoverer search engine (v2.4.1.15). Tandem mass spectra were searched against the Mus musculus SwissProt database (17,063 entries), concatenated with a reverse decoy database. Trypsin (full) was specified as the cleavage enzyme, allowing up to two missing cleavages. The mass tolerance for precursor ions was set to 10 ppm in the first search and the mass tolerance for fragment ions was set to 0.02 Da. Carbamidomethyl on Cys was specified as the fixed modification, and acetylation on protein N-terminal, oxidation on Met, deamidation (NQ), and phosphorylation on Ser, Thr, and Tyr were specified as variable modifications. The FDR was adjusted to <1%. All the other parameters in the Proteome Discoverer were set to default values.

The raw LC-MS datasets were first searched against the database and converted into matrices containing the intensity of peptides across samples. The relative quantitative and statistical values of each modified peptide were then calculated based on this intensity information by the following steps:

1. Peptides with a modification pattern described as “positions not distinguishable” and Quan Info marked by “NoQuanValues” were removed from all peptide datasets.
2. The intensities of modified peptides (I) were centralized and transformed into relative quantitative values (R) of modified peptides in each sample using the following formula: Rij = Iij / Mean(Ij). Here, i denotes the sample and j denotes the modified peptide. If both proteomics and post-translational modification profiling were conducted on the same cohort, the relative quantitative value of the modified peptide was typically divided by the relative quantitative value of the corresponding protein to remove the influence from protein expression of modifications.
3. Average relative quantitative values were calculated for each group, and the log2 ratio values of *Cdk*13 KD compared with control groups were also calculated.
4. P-values were calculated with log2 relative quantitative values using a t-test. The modified sites with |fold change | > 1.5 and P < 0.1 were regarded as differentially expressed phosphopeptides. Gene Ontology enrichment analysis was carried out using a method similar to that used for RNA-seq analysis.

### Software

HISAT2 v2.1.0

featureCounts v1.6.2

R v4.0.3

DESeq2 v1.30.0

clusterProfiler v3.18.0

org.Mm.eg.db v3.12.0

Bowtie2 v2.4.2

samtools v1.3.1

sambamba v0.8.1

macs2 v2.2.7.1

deepTools v3.5.1

ChIPseeker v1.22.1

TxDb.Mmusculus.UCSC.mm10.knownGene v3.10.0

ggplot2 v3.3.5

pheatmap v1.0.12

UCSC genome browser http://genome.ucsc.edu

GraphPad Prism 8 version 8.3.0(538)

mm10-blacklist.v2

Cytoscape 3.8.0

MoMo Version 5.4.1 https://meme-suite.org/meme/tools/momo

## Data availability

All genomic data are being made accessible at the Gene Expression Omnibus (GEO) database repository. ChIP-seq: GSE190679; RNA-seq: GSE190781, GSE190692, GSE190683. Phosphoproteomic data are being made accessible at the PRoteomics IDEntifications database (PRIDE) (https://www.ebi.ac.uk/pride) repository PXD030231.

## Acknowledgements

We thank Professor An-Dong Qiu for generously providing us with cre mouse strains as gifts and for his helpful suggestions during the study. Professor Ci-Zhong Jiang and his group members generously helped with our bioinformatic analysis. We also thank Da-De Song and Dr Feng Sun for their technical assistance. We would also like to thank Professor Zhi-Yong Mao for his helpful suggestions during the study.

## Author contributions

C.W., W.Z., X.L.C., W.F.W., X.B.G., J.F.C., Y.B.L., K.L., Y.Y.J., X.Y.L. and Y.T.L. conducted the experiments; Y.H.L. performed the bioinformatic analysis; C.W. performed the ChIP-seq preparation, ChIP‒qPCR, and IP experiments; W.Z. and Y.Y.J. helped with the KO mice and immunofluorescence (IF) experiments; W.F.W., X.B.G. and K.L. helped with cell line preparation and IF staining; X.L.C. and Y.B.L. helped with myocardial-specific deletion work in mice; J.F.C. produced the mutant plasmid and performed protein interaction analysis; and W.F.W. assisted in preparing the graphics for the manuscript.

## Additional information

Correspondence and requests for materials should be addressed to Fang-Lin Sun

---

**Supplementary Fig. 1.**
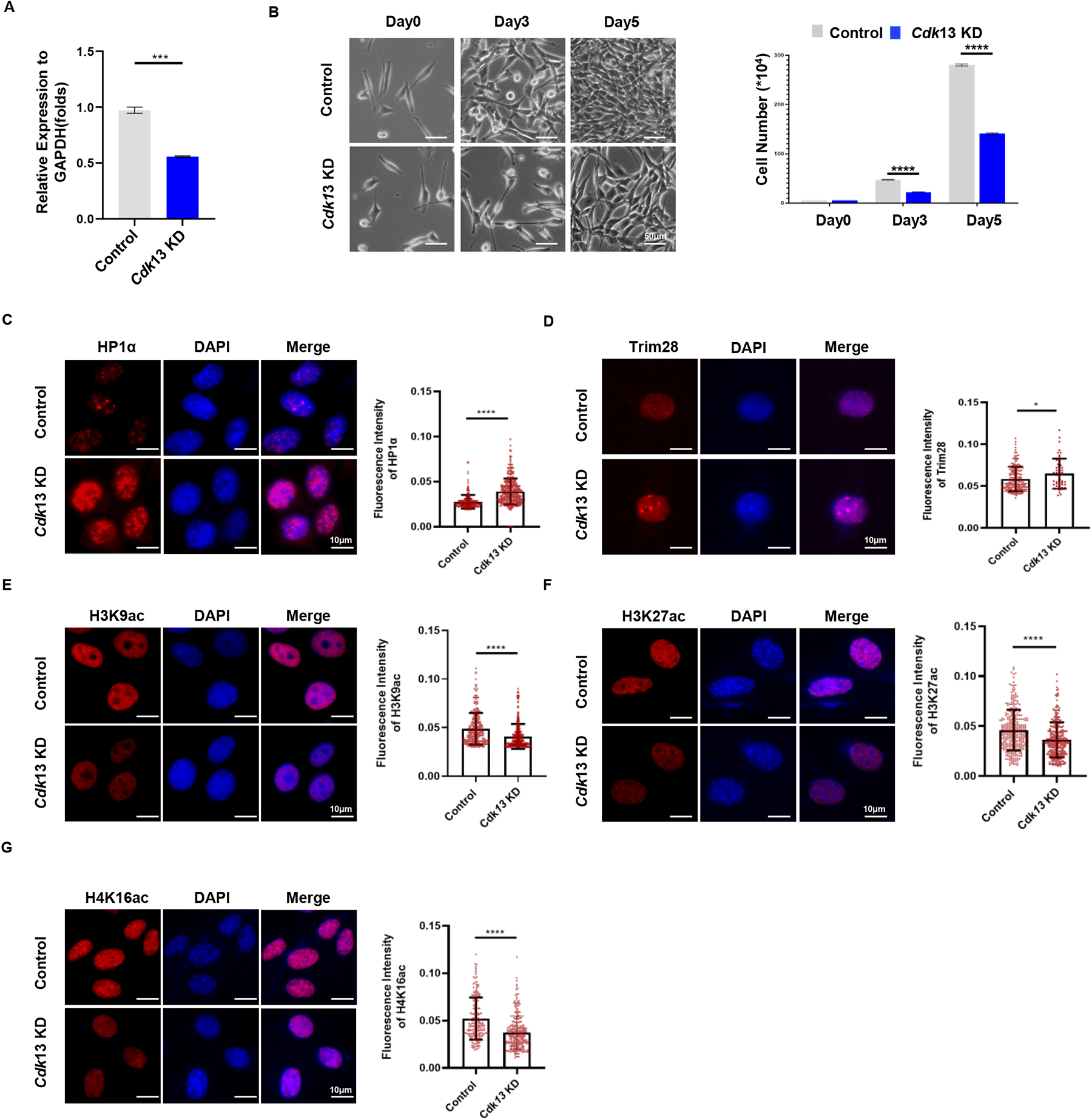
Depletion of *Cdk*13 inhibits cell proliferation and markers of active chromatin. **A***, Cdk*13 was efficiently knockdown (KD) by shRNA in MEFs. RT-qPCR was performed to examine the expression of *Cdk*13. The data are presented as the means ± SEMs. \*\*\**P* < 0.001. t-test. **B**, *Cdk*13 KD compromised cell proliferation. The images of cells on day 0, day 3, and day 5 (left panel), and the quantification of cell numbers (right panel) are shown. The data are presented as the means ± SEMs. \*\*\*\**P* < 0.0001, t-test. **C-G**, The left panel shows representative immunofluorescence staining for HP1α (c), Trim28 (d), H3K9ac (e), H3K27ac (f), and H4K16ac (g) in control and *Cdk*13 KD mouse embryonic fibroblasts (MEFs) with lentivirus shRNA. The right panel shows the quantification of fluorescence intensity of the corresponding images. Each point represents a cell, and more than 300 cells per condition were analyzed. The data are presented as the means ± SEMs. *P < 0.05, ****P < 0.0001, t-test.

**Supplementary Fig. 2.**
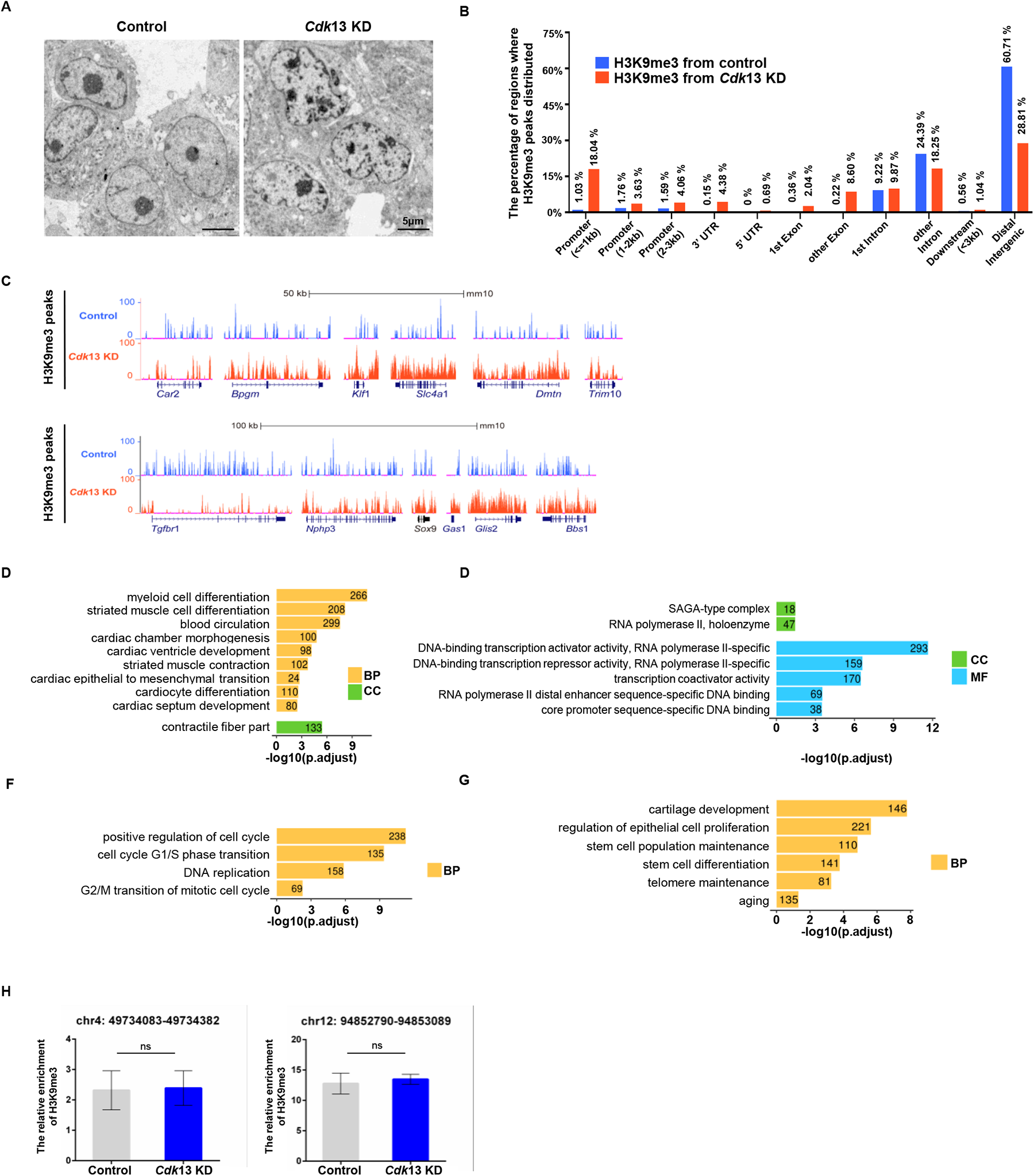
Depletion of *Cdk*13 promotes H3Kme3-heterochromatin spreading. **A**, Representative transmission electron microscopy images show heterochromatin in control and MEFs with *Cdk*13 KD. n = 15 nuclei. **B**, The relative percentage of H3K9me3 in the promoter region, exon region, 3’ UTR and distal intergenic regions of control and *Cdk*13 KD MEFs. The bar plot shows the H3K9me3 peak percentage of each region. The blue bar represents the control group, and the red bar represents the *Cdk*13 KD group. **C**, The representative genes show the distribution of H3K9me3 peaks. The views of H3K9me3 in the UCSC genome is shown. The blue represents the control group, and the red represents the *Cdk*13 KD group. **D**-**G**, Selected Gene Ontology (GO) terms enriched with H3K9me3 in the promoter after *Cdk*13 KD. The column in yellow, green and blue represents genes involved in biological process (BP), cellular component (CC), and molecular function (MF). The x-axis represents the -log10 (adjusted P value), and the numbers on the bars are the numbers of genes enriched for each term. **H**, ChIP-qPCR results verified the intergenic region where H3K9me3 showed no difference.

**Supplementary Fig. 3.**
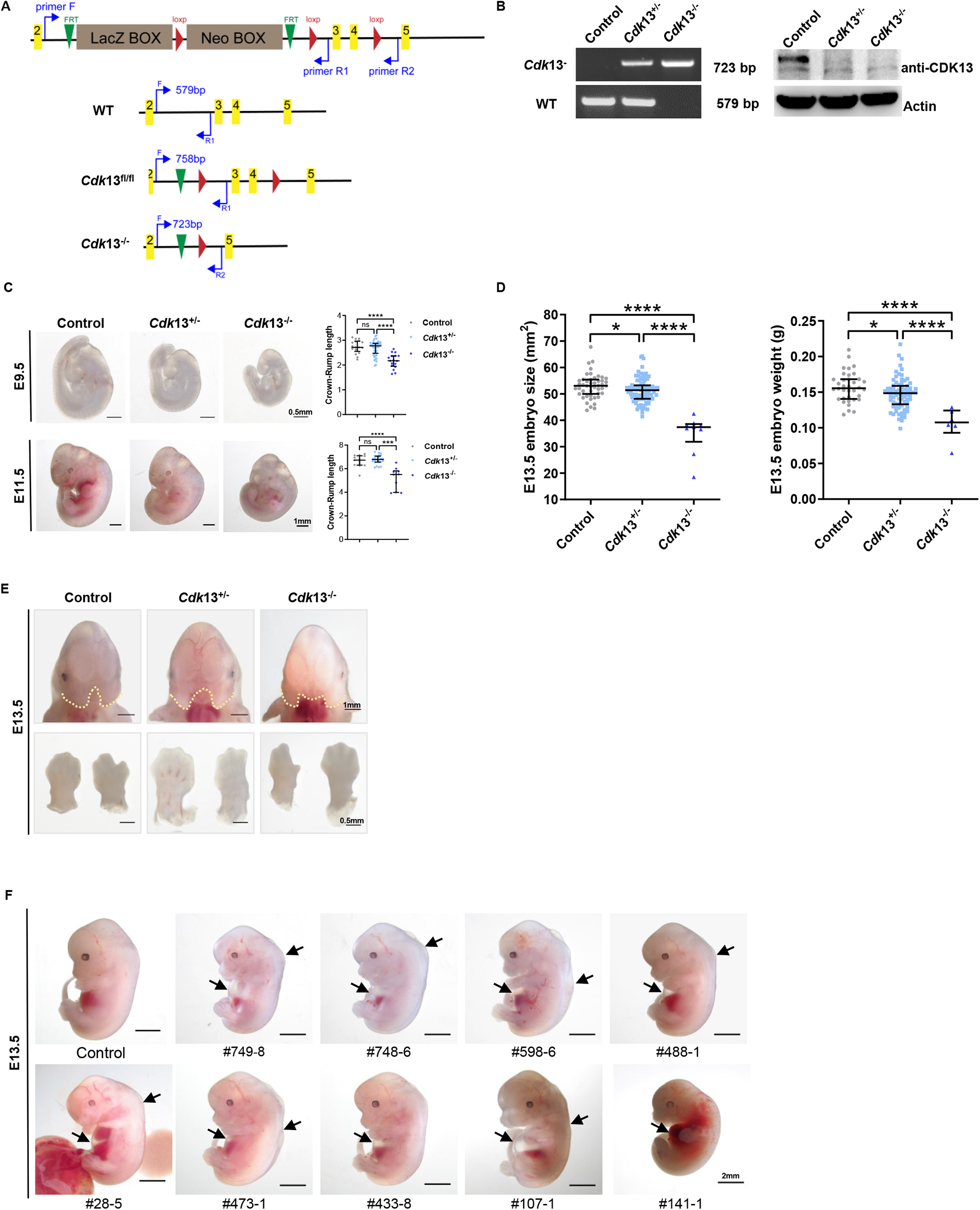
*Cdk*13 knockout in mice causes embryonic morphology defects. **A**, Schematic illustration of the constructs used to generate *Cdk*13 knockout (*Cdk*13^-/-^) mice and PCR primers monitoring the deletion efficiency of *Cdk*13^-/-^mice. **B**, The deletion efficiency of *Cdk*13^-/-^ was confirmed by PCR in the left panel and Western blot analysis in the right panel. E13.5 embryo tail tissue DNA was used for PCR. E13.5 mouse embryonic brain tissue was used in the Western blotting to detect Cdk13 protein expression. Actin was used as an internal control. **C**, *Cdk*13 knockout (*Cdk*13^-/-^) resulted in smaller size embryos at E9.5 and E11.5 than controls. The right panel shows the quantification of the Crown-Rump length of E9.5 and E11.5 embryos. The number of embryos at E9.5 was Control, n=17; *Cdk*13^+/-^, n=44; *Cdk*13^-/-^, n=17. The number of embryos at E11.5 was control, n=13; *Cdk*13^+/-^, n=24; *Cdk*13^-/-^, n=10, with a scale bar of 0.5mm. The data are presented as the means ± SEMs. \*\*\**P* < 0.001, \*\*\*\**P* < 0.0001, t-test. **D** Quantification of the size and weight of E13.5 embryos. The number of embryos: control, n=36; *Cdk*13^+/-^, n=74; *Cdk*13^-/-^, n=6. **E**, *Cdk*13^-/-^ resulted in craniofacial defects (upper panel) and digital anomalies (lower panel) in mouse E13.5 embryos. The scale bar was 0.5mm for digital anomalies and 1mm for craniofacial defects. **F**, *Cdk*13 knockout resulted in pleural/back effusion and smaller size of mouse embryos at E13.5. The unique code number of each fetus is also indicated.

**Supplementary Fig. 4.**
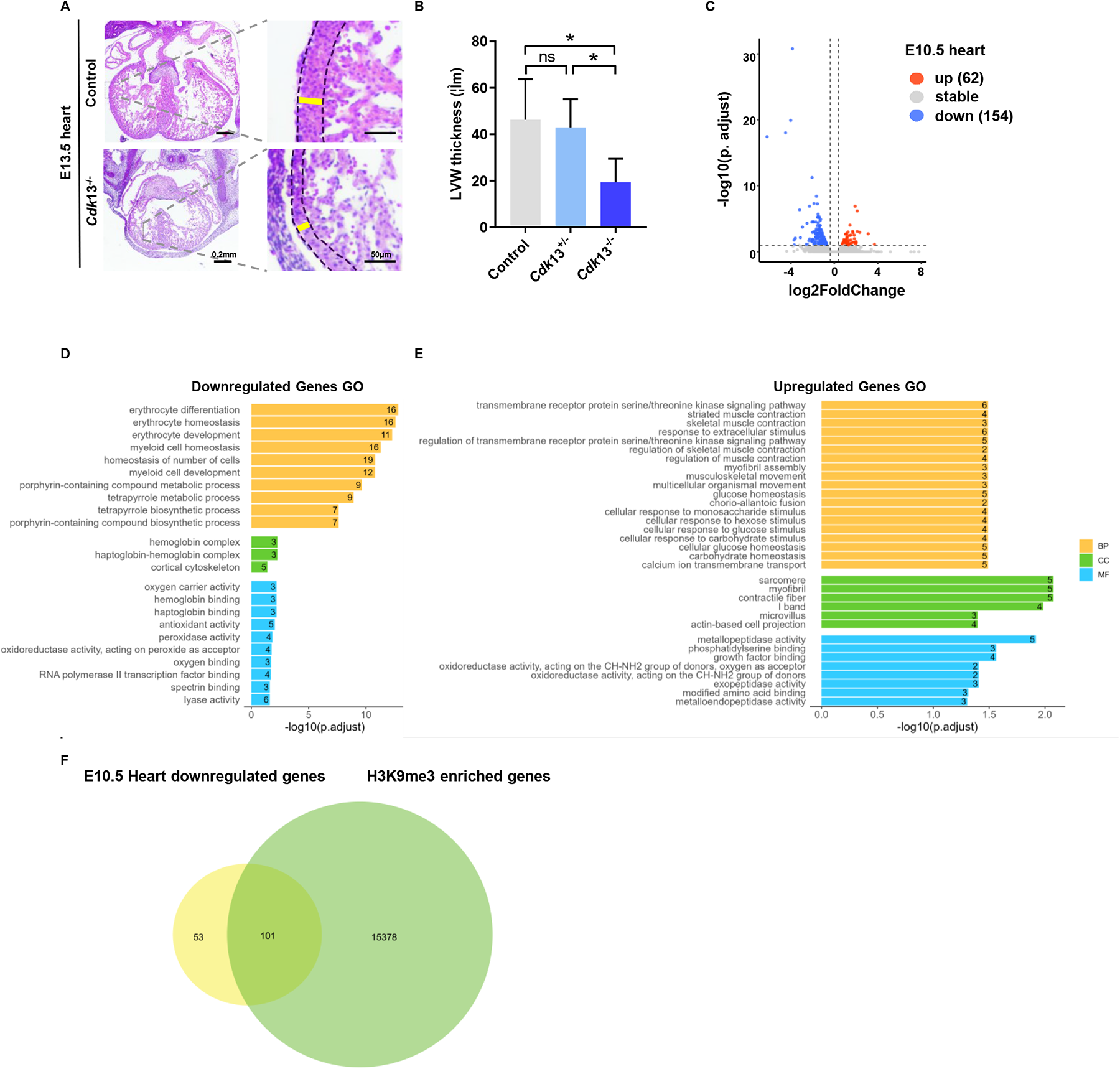
*Cdk*13 deletion causes embryo heart defects and downregulation of genes. **A**, *Cdk*13 knockout resulted in a defective interventricular septum and decreased thickness of the ventricular walls in E13.5 embryos. The scale bar was 200μm (left) and 50μm (right). **B**, Quantification of the thickness of the ventricular wall using E13.5 *Cdk*13^-/-^ mutant and control heart tissues. The data are presented as means ± SEMs. N=3, \**P* < 0.05, t-test. **C**, The volcano plot of RNA sequencing data shows differentially expressed genes (DEGs; fold change>1.3 and adjusted *P* < 0.1) in E10.5 heart. The number of downregulated genes is 2.5 folds of the number of upregulated genes. The red points represent upregulated genes, and the blue points represent downregulated genes; the dotted line indicates the threshold of the fold change and adjusted *P* value. **D**, **E**, Gene Ontology analysis of the altered expression of genes from RNA-seq. **F**, Two-thirds of downregulated genes in E10.5 hearts from RNA-seq overlapped with the corresponding genes showing increased H3K9me3 in *Cdk*13 KD MEFs (Fig. 1c).

**Supplementary Fig. 5.**
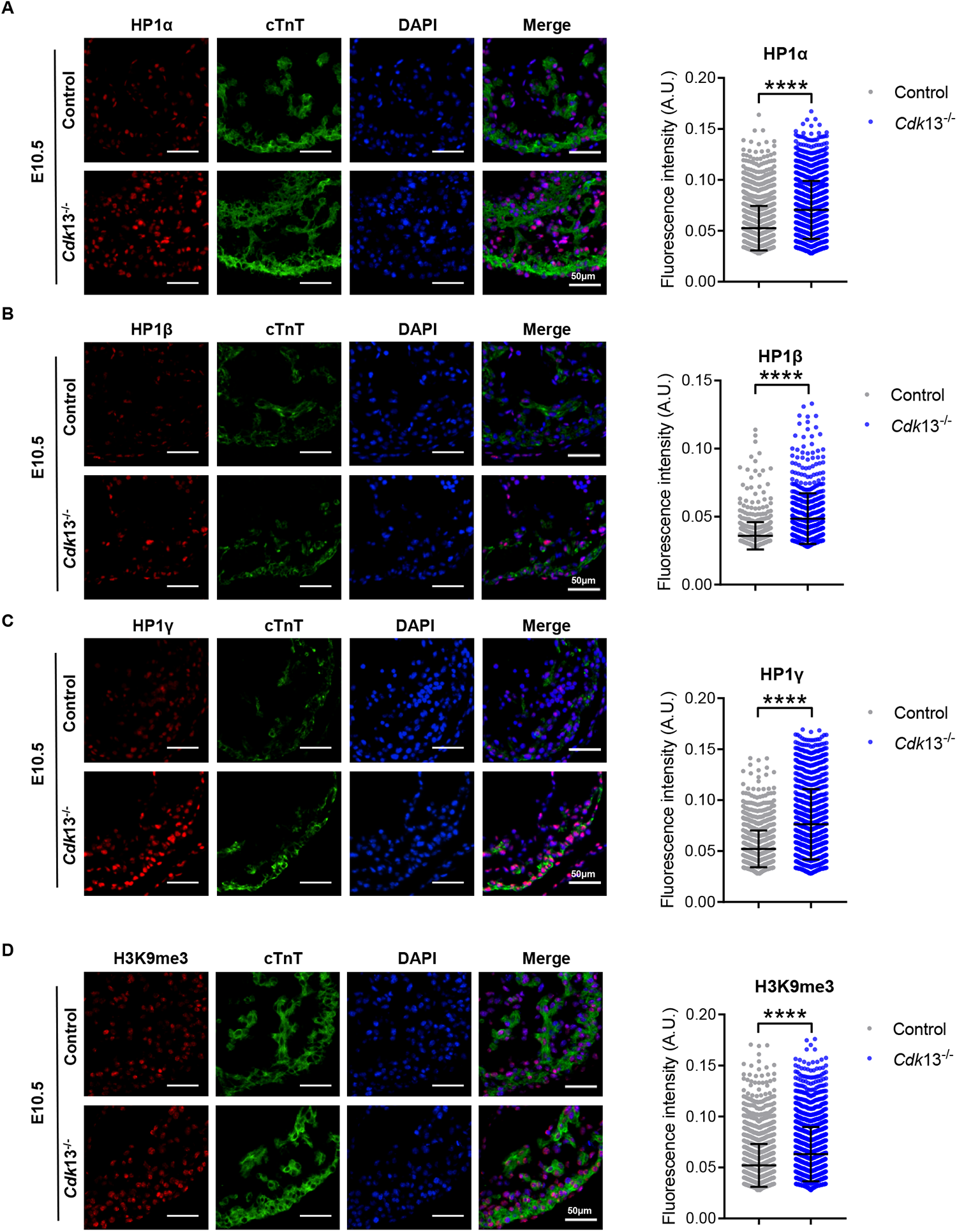
*Cdk*13 deletion increases heterochromatinization in E10.5 mouse embryonic heart tissues. The left panel shows immunofluorescence staining performed using antibodies against markers of heterochromatin, HP1α (**A)**, HP1β (**B)**, HP1γ (**C)**, and H3K9me3 (**D)**, which are labeled in red. Cardiomyocytes labeled with cTnT are green, and DAPI is shown in blue. All images show the left ventricle position. The scale bar was 50μm. The right panel shows the fluorescence intensity of single cell, and more than 200 cells per condition were analyzed (**A**-**D**). A. U, arbitrary unit, N = 3/4/5, t-test. ****P < 0.0001.

**Supplementary Fig. 6.**
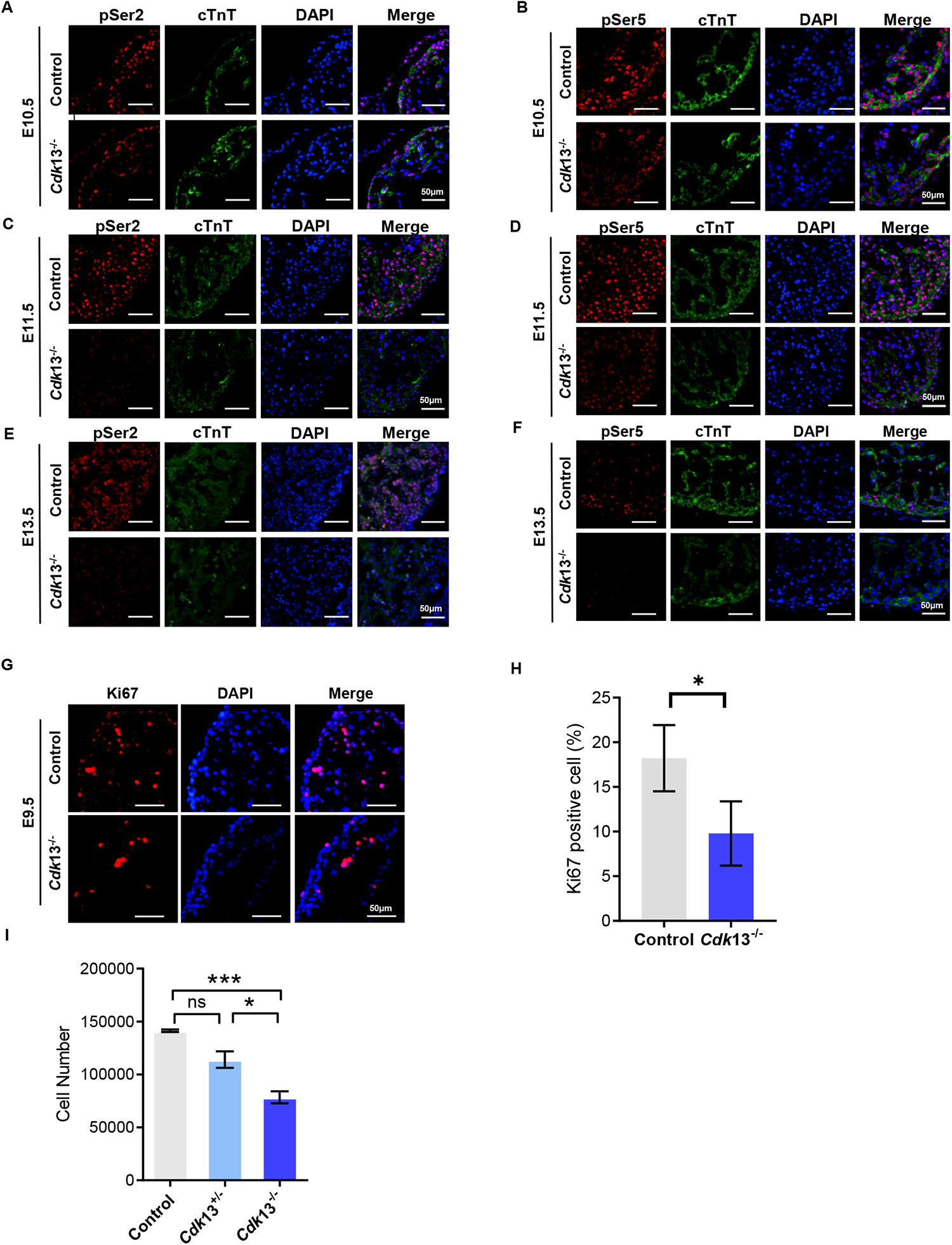
*Cdk*13 deletion decreases phosphorylation of RNA polymerase II and cell proliferation. **A-F**, Immunofluorescence staining was performed using antibodies against RNA polymerase II pSer2 (left) and pSer5 (right) in E10.5 (**A, B**), E11.5 (**C, D**) and E13.5 (**E, F**) embryonic heart tissues. All images show the left ventricle position. The scale bar was 50 μm. **G**, Immunofluorescence staining was performed using anti-Ki67 in E9.5 mouse embryonic heart tissues. The scale bar was 50 μm. **H**, Quantification of Ki67-positive cells. N=3 *P <0.1, t-test (**I**) The total number of cells was decreased in E13.5 *Cdk*13^-/-^ mouse embryos. The analyzed data are presented as means ± SEMs. N=3 \**P* < 0.05, \*\*\**P* < 0.001, t-test.

**Supplementary Fig. 7.**
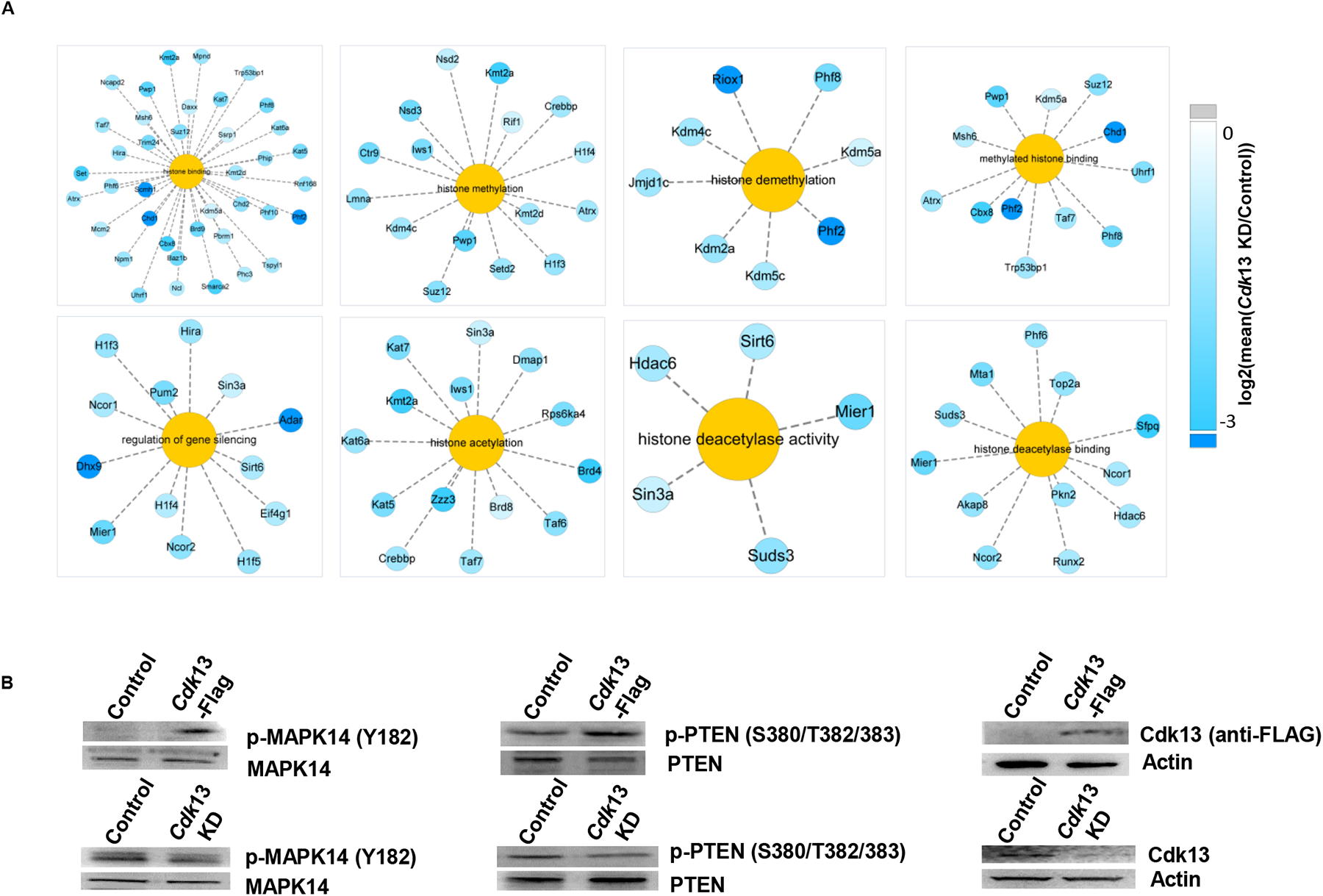
Gene Ontology enrichment analysis and validation of Cdk13-dependent protein phosphorylation regulated by Cdk13. **A**, simple network plot shows the selected Gene Ontology terms and related proteins with decreased phosphorylation upon *Cdk*13 KD, including histone binding, histone methylation and histone demethylation, histone acetylation and deacetylation. The colours of the protein nodes represent the log2 (mean fold change) values. **B**, The Western blotting analysis shows that the phosphorylations of MAPK14 and PTEN were regulated by Cdk13 in a dosage-dependent manner. The top panel shows the increased phosphorylation of p-MAPK14(Y182) and p-PTEN(S380/T382/383) in the presence of Cdk13 overexpression. The bottom panel shows the decreased phosphorylation of p-MAPK14(Y182) and p-PTEN(S380/T382/383) after depletion of *Cdk*13. The overexpression of Cdk13 was monitored using anti-FLAG antibody, and the Cdk13 depletion efficiency was analyzed using the anti-CDK13 antibody.

**Supplementary Fig. 8.**
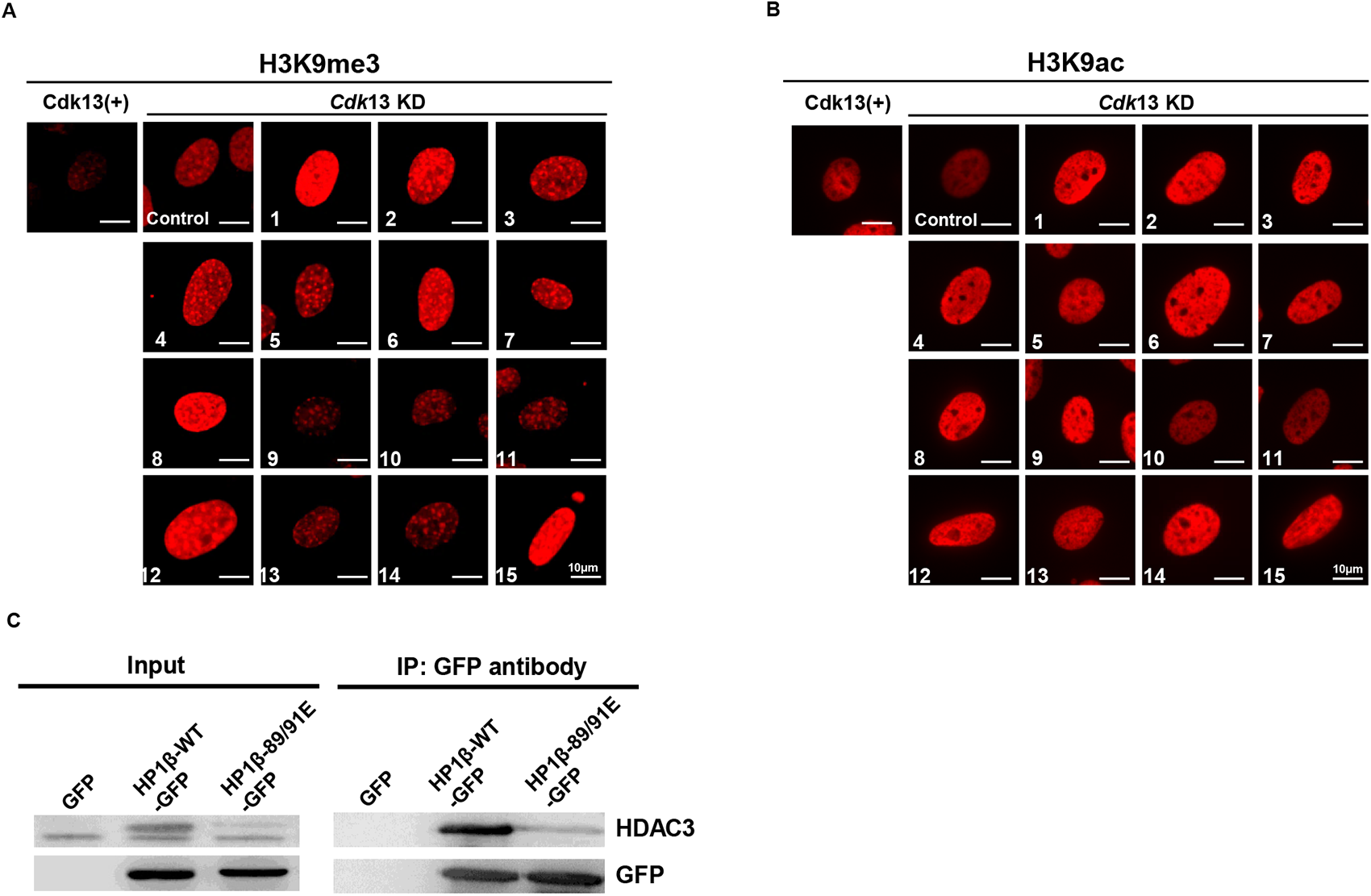
A screen of small molecules antagonizing H3K9me3-heterochromatin. **A, B**, Representative immunofluorescence images for H3K9me3 and H3K9ac (red) in the presence of small molecules. **C**, IP experiments show the interaction between wild-type HP1β with HDAC3.

**Supplementary Fig. 9.**
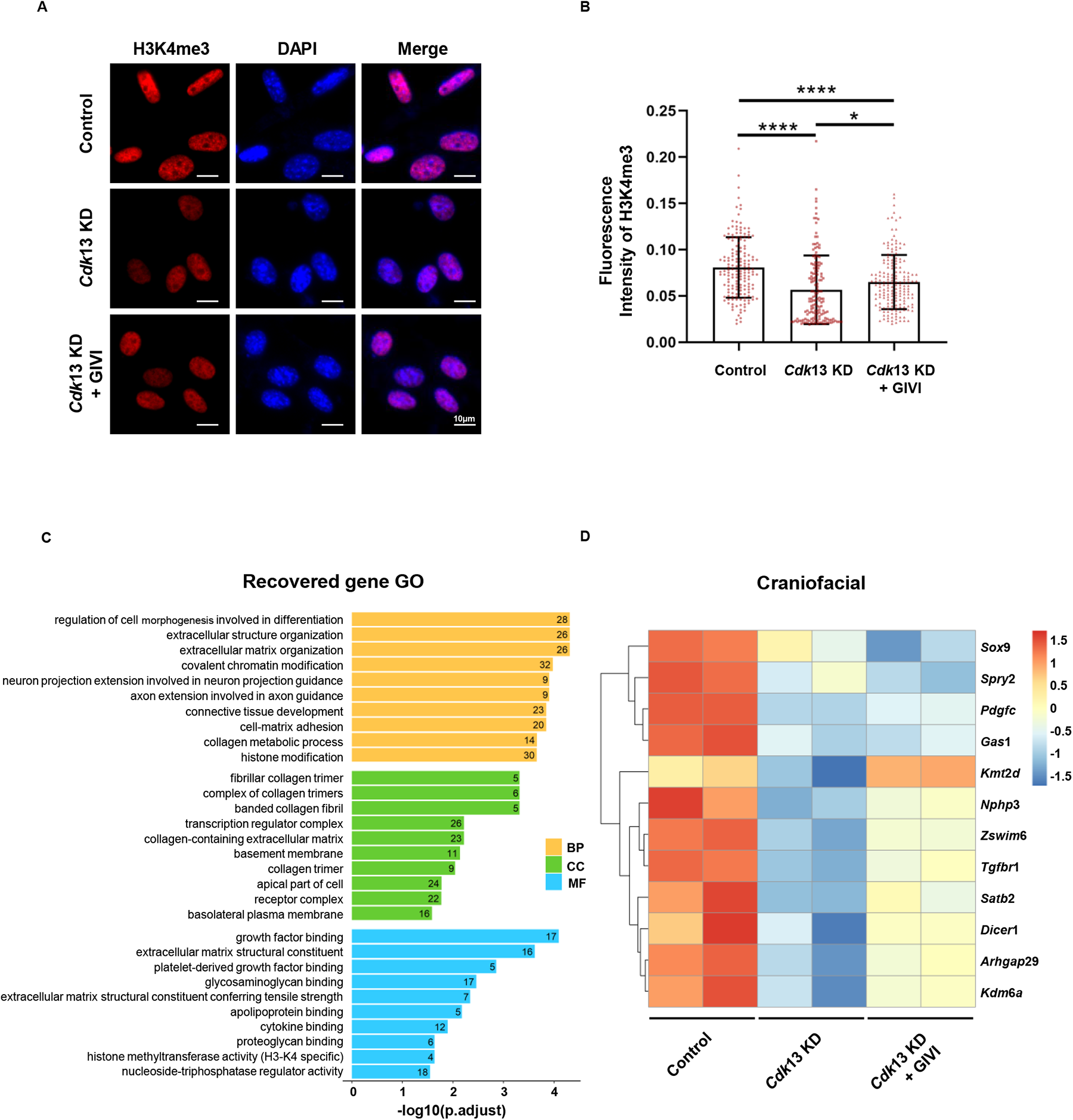
GIVI prevents heterochromatinization to facilitate transcriptional activation. **A**, GIVI partially recovers the intensity of H3K4me3 (red) caused by *Cdk*13 KD in MEFs. **B**, The quantification of H3K4me3 intensity is shown. Each point represents a cell, and more than 300 cells were analysed under the same conditions. The data are presented as means ± SEMs. \**P* < 0.05, \*\*\*\**P* < 0.0001, t-test. **C**, Gene Ontology enrichment analysis of genes rescued by GIVI. Yellow, green and blue columns represent genes involved in biological process (BP), cellular component (CC) and molecular function (MF), respectively. **D**, A representative heatmap shows the RNA sequencing analysis of the control, *Cdk*13 KD and *Cdk*13 KD + GIVI groups. A number of differentially expressed genes (DEGs) related to craniofacial development are shown.

**Supplementary Fig. 10.**
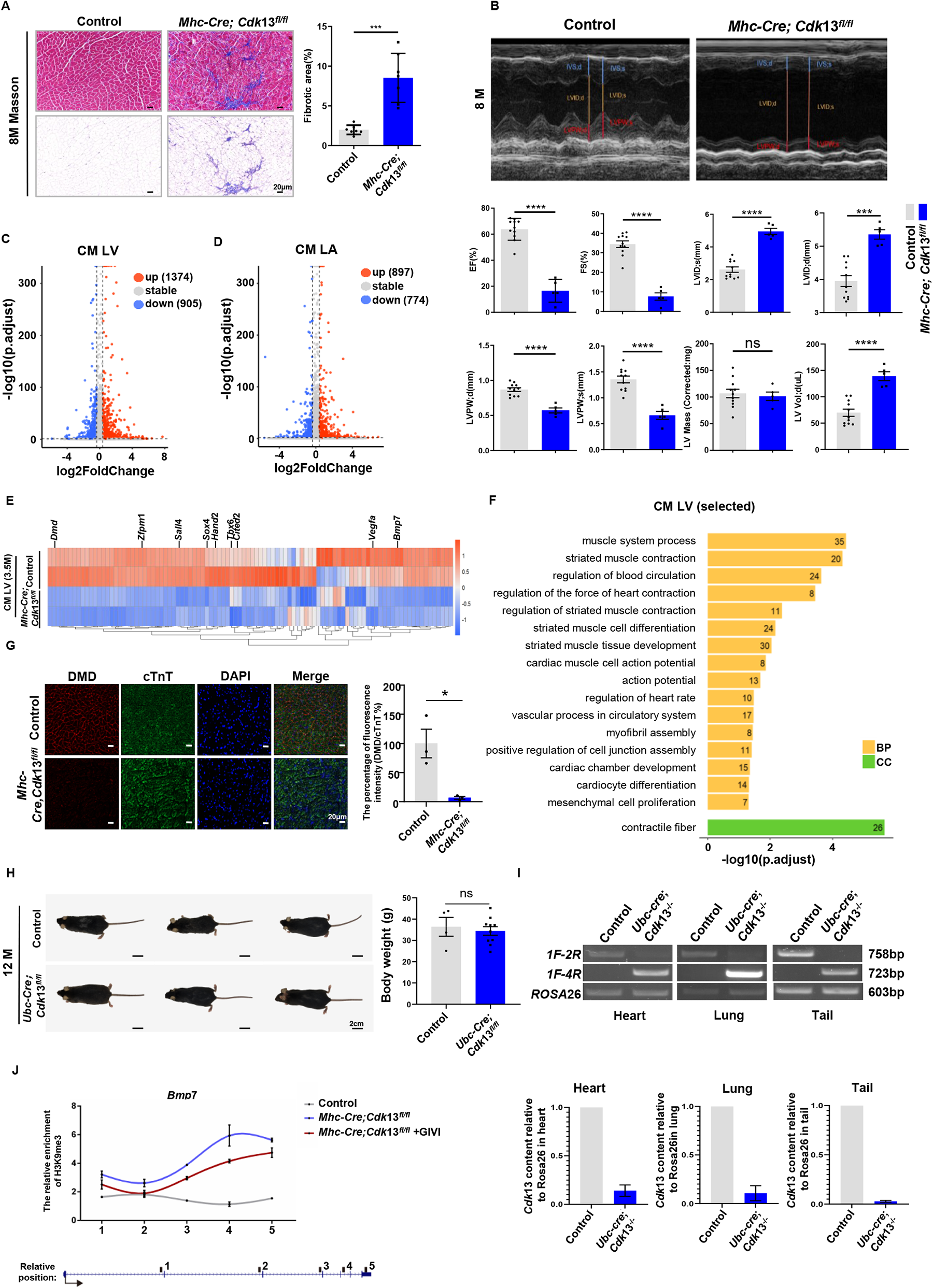
Ph enotypic and transcriptional analysis of cardiomyocyte-specific deletion of *Cdk*13 in mice and *Cdk*13 deletion-induced H3K9me3 changes in prenatal mouse hearts. **A**, Fibrosis significantly accumulated in the hearts of *Mhc-cre;Cdk*13*^fl/f^*^l^ mice. The left panel shows the heart layers of 8-month-old control and *Mhc-Cre;Cdk*13*^fl/fl^*mice using Masson’s staining to indicate the pathological fibrosis process. The right panel shows the quantification analysis of cardiac fibrosis area. The fibrosis area (blue) in the heart tissue was assessed in Photoshop, and the area was calculated and then divided by the total area to obtain the fibrosis area; t-test, \*\*\**P* < 0.001, *n* = 7. **B**, Echocardiographic images showed the functional defects by *Cdk*13 deletion in heart. Eleven control and five knockout mice were analysed. IVS, interventricular septum; LVID, left ventricular internal diameter; LVPW, left ventricular posterior wall. EF, ejection fraction; FS, fractional shortening; LVPW;s, left ventricular posterior wall, systolic; LVPW;d, left ventricular posterior wall, diastolic. T-test, **N=**11(Control)/5(*Mhc-Cre*; *Cdk*13^fl/fl^), ns, not significant; \*\*\**P* < 0.001; \*\*\*\**P* < 0.0001, t-test. **C,D**, Volcano plot of DEGs (fold change >1.3 and adjusted *P* < 0.1) in the left ventricle (LV) and left atrium (LA) of 3.5-month-old mice. CM: cardiac muscle. Red points indicate upregulated genes, blue points indicate downregulated genes, and the dotted line represents the threshold of the fold change and adjusted *P* value. **E**, Heatmap of the downregulated genes related to selected Gene Ontology terms, with a list of genes essential for heart development displayed above. **F**, Selected terms from Gene Ontology enrichment analysis of downregulated genes in *Cdk*13 mutant. Yellow represents biological process (BP); green represents cellular component (CC). The x-axis represents the –log10 (adjusted *P* value) value, and the numbers on the bars indicate the numbers of genes enriched for each term. **G**, Cardiomyocyte-specific deletion of *Cdk*13 resulted in a significant decrease in dystrophic protein (Dmd) expression. The left panel shows fluorescence staining results using antibodies against Dmd (red) or cTnT (green) of control or *Mhc-Cre;Cdk*13*^fl/fl^* hearts of 7-month-old mice. The right panel shows the quantification of the relative intensity of Dmd. The data are presented as means ± SEMs. \**P* < 0.05, t-test. n=3. **H**, The left panel represents the body panorama of Control or Ubc-Cre; *Cdk*13*^fl/fl^*, the right panel shows statics of body weight(g) in Control or *Ubc-creERT2*; *Cdk*13*^flfl^* 12-month-old mice. The data are presented as means ± SEMs. \**P* < 0.05, t-test. **I**, *Cdk*13 deletion efficiency verification. The top panel shows the PCR verification results of *Cdk*13 deletion efficiency in *Ubc*-*creERT2*; *Cdk*13*^fl/fl^*mice heart(right), lung(middle), and tail(left). The lower panel shows the quantification of the 758bp PCR products. *Rosa*26 was used as the internal control. J. ChIP-qPCR analysis showed that the enrichment of H3K9me3 in *Cdk*13 knockout embryos was decreased in the presence of GIVI. The primers were designed to amplify the defined regions in the *Bmp*7.

**Supplementary Table. 1.** The genotype and number of surviving *Cdk*13^-/-^ embryos at different stages.

| Stage | *Cdk13*<sup>+/+</sup>embryos<br>(expected 25%) | *Cdk13*<sup>+/-</sup>embryos<br>(expected 50%) | *Cdk13*<sup>-/-</sup>embryos<br>(expected 25%) | Empty yolk<br>sac/dead | Total<br>number of<br>embryos |
| --- | --- | --- | --- | --- | --- |
| E9.5 | 23 (20.91%) | 54 (49.09%) | 21 (19.09%) | 12 (10.91%) | 110 |
| E10.5 | 48 (30.97%) | 58 (37.42%) | 37 (23.87%) | 12 (7.74%) | 155 |
| E11.5 | 15 (23.08%) | 29 (44.62%) | 11 (16.92%) | 10 (15.38%) | 65 |
| E13.5 | 75 (26.04%) | 143 (49.65%) | 14 (4.86%) | 56 (19.44%) | 288 |
| E15.5 | 8 (20.00%) | 22 (55.00%) | 0 (0.00%) | 10 (25.00%) | 40 |

**Supplementary Table. 2.** Phenotype of E13.5 Cdk13^-/-^ embryos.

| Number | Genotype | Phenotype of E13.5 *Cdk13*<sup>-/-</sup> embryos |
| --- | --- | --- |
| #28-5 | *Cdk13*<sup>-/-</sup> | Craniofacial defects, digital anomalies (front left), pleural effusion, back effusion |
| #28-6 | *Cdk13*<sup>-/-</sup> | Craniofacial defects, digital anomalies (front right), pleural effusion, back effusion |
| #72-1 | *Cdk13*<sup>-/-</sup> | Craniofacial defects, digital anomalies (front right), pleural effusion, back effusion |
| #107-1 | *Cdk13*<sup>-/-</sup> | Smaller size, Craniofacial defects, digital anomalies (front right), pleural effusion, back effusion |
| #135-1 | *Cdk13*<sup>-/-</sup> | Smaller size, craniofacial defects, digital anomalies (front left), pleural effusion, back effusion |
| #141-1 | *Cdk13*<sup>-/-</sup> | Craniofacial defects, digital anomalies, pleural effusion, defective neural tube |
| #433-8 | *Cdk13*<sup>-/-</sup> | Smaller size, digital anomalies (front right), pleural effusion |
| #473-1 | *Cdk13*<sup>-/-</sup> | Smaller size, craniofacial defects, digital anomalies (front right), pleural effusion, back effusion |
| #488-1 | *Cdk13*<sup>-/-</sup> | Smaller size, craniofacial defects, digital anomalies (front right), pleural effusion, back effusion |
| #598-5 | *Cdk13*<sup>-/-</sup> | Smaller size, pleural effusion, back effusion. |
| #748-6 | *Cdk13*<sup>-/-</sup> | Smaller size, craniofacial defects, pleural effusion, back effusion, digital anomalies (front left) |
| #749-8 | *Cdk13*<sup>-/-</sup> | Smaller size, craniofacial defects, pleural effusion, back effusion, digital anomalies (front left and front right) |
| #20-1 | *Cdk13*<sup>-/-</sup> | No obvious defects |
| #598-1 | *Cdk13*<sup>-/-</sup> | No obvious defects |
| #26-4 | *Cdk13*<sup>-/-</sup> | Prenatal lethality, craniofacial defects |

**Supplementary Table 3.** Specific information and working concentrations of 15 small molecule inhibitors used for screening.

|  | Catelog | Name | Molecular weight | Cas number | concentration |
| --- | --- | --- | --- | --- | --- |
| 1 | S1095 | Dacinostat (LBH589) | 379.459 | 404951-53-7 | 0. 2uM |
| 2 | S1422 | Droxinostat(LAQ824) | 243.69 | 99873-43-5 | 10uM |
| 3 | S2012 | PCI-34051 | 296.32 | 950762-95-5 | 5uM |
| 4 | S2818 | Tacedinaline(CI994) | 269.3 | 112522-64-2 | 5uM |
| 5 | S7229 | RGFP966 | 362.4 | 1396841-57-8 | 2uM |
| 6 | S7473 | Nexturastat A | 341.4 | 1403783-31-2 | 10uM |
| 7 | S8001 | Ricolinostat(ACY-1215) | 433.5 | 1316214-52-4 | 2uM |
| 8 | S8043 | Scriptaid | 326.35 | 287383-59-9 | 5uM |
| 9 | S2170 | Givinostat (ITF2357) Givi | 475.97 | 732302-99-7 | 2.5nM |
| 10 | S7237 | OG-L002 | 225.29 | 1357302-64-7 | 2uM |
| 11 | S7574 | GSK-LSD1 2HCl | 289.24 | 1431368-48-7 | 2uM |
| 12 | S7680 | SP2509 | 437.9 | 1423715-09-6 | 2uM |
| 13 | S7796 | GSK2879552 2HCl | 437.4 | 1401966-69-5 (free pase) | 2uM |
| 14 | S7572 | A-366 | 329.44 | 1527503-11-2 | 1uM |
| 15 | S7591 | BRD4770 | 413.47 | 1374601-40-7 | 10uM |

